# Reverse plasticity underlies rapid evolution by clonal selection within populations of fibroblasts propagated on a novel soft substrate

**DOI:** 10.1101/2020.04.26.061911

**Authors:** Purboja Purkayastha, Kavya Pendyala, Ayush S. Saxena, Hesamedin Hakimjavadi, Srikar Chamala, Purushottam Dixit, Charles F. Baer, Tanmay P. Lele

## Abstract

Mechanical properties such as substrate stiffness are a ubiquitous feature of a cell’s environment. Many types of animal cells exhibit canonical phenotypic plasticity when grown on substrates of differing stiffness, *in vitro* and *in vivo*. Whether such plasticity is a multivariate optimum due to hundreds of millions of years of animal evolution, or instead is a compromise between conflicting selective demands, is unknown. We addressed these questions by means of experimental evolution of populations of mouse fibroblasts propagated for ~90 cell generations on soft or stiff substrates. The ancestral cells grow twice as fast on stiff substrate as on soft substrate and exhibit the canonical phenotypic plasticity. Soft-selected lines derived from a genetically diverse ancestral population increased growth rate on soft substrate to the ancestral level on stiff substrate and evolved the same multivariate phenotype. The pattern of plasticity in the soft-selected lines was opposite of the ancestral pattern, suggesting that reverse plasticity underlies the observed rapid evolution. Conversely, growth rate and phenotypes did not change in selected lines derived from clonal cells. Overall, our results suggest that the changes were the result of genetic evolution and not phenotypic plasticity per se. Whole-transcriptome analysis revealed consistent differentiation between ancestral and soft-selected populations, and that both emergent phenotypes and gene expression tended to revert in the soft-selected lines. However, the selected populations appear to have achieved the same phenotypic outcome by means of at least two distinct transcriptional architectures related to mechano-transduction and proliferation.

## Introduction

Mechanical properties such as pressure, viscosity and substrate stiffness are inherent components of a cell’s environment. This is true for both unicellular microbes and for somatic cells in a multicellular organism. Just as temperature and ambient chemistry often vary over the course of the life of a cell and its recent descendants, mechanical properties may be similarly variable. Therefore, we expect that mechanical properties of a cell’s environment constitute a significant agent of natural selection; but this has never been investigated before to our knowledge.

Depending on the time scale and predictability of the environmental variation, natural selection may favor phenotypic plasticity, wherein an individual of a given genotype develops a different set of traits depending on the environmental context (Ghalambor et al. 2007; Price et al. 2003). Evolutionary biologists are used to thinking about phenotypic plasticity in the context of autonomous individual organisms, but the same principle applies to somatic cells within a developing multicellular organism.

In fact, animal somatic cells exhibit remarkably strong and consistent plasticity in response to the mechanical environment. For instance, mammary epithelial cells proliferate faster in the terminal end buds of developing mammary glands in mice where the extracellular matrix (ECM) is stiff, while they proliferate slower in the duct where the matrix is soft. This differential proliferation results in elongated and laterally branched structures (Gjorevski and Nelson 2011; Nelson and Gleghorn 2012). Similar branching occurs in capillary networks as a result of endothelial cellular response to spatial gradients in matrix stiffness that drive tissue patterning during angiogenesis (Huang and Ingber 1999; Ingber 2002). Likewise, mesodermal stiffness gradients bias collective cell migration from soft to stiff regions to provide shape to the early limb bud in developing mouse embryos (Zhu et al. 2020). Separately from development, stiffness gradients guide migration of fibroblasts towards the wound during wound healing (Lancerotto and Orgill 2014). In addition to development and normal biological processes, cellular responses to changes in matrix stiffness are associated with pathologies including cancer (Chin et al. 2016). For example, tumor tissue with elevated stromal stiffness promotes malignant cancer cell phenotypes and functions like increased proliferation, altered tissue polarity and a lack of lumen in breast cancer, prostate cancer and melanoma (Levental et al. 2009; Paszek et al. 2005; Wang et al. 2014; Weder et al. 2014)

Consistent with such *in vivo* observations, different somatic cell types display consistent differences when cultured on soft substrates compared to stiff substrates in the rate of proliferation, apoptosis, differentiation, cell spreading and migration (Janmey et al. 2020; Mih et al. 2011; Peyton and Putnam 2005). Mechanical stiffness also affects intracellular features, including cytoskeletal organization, nuclear shape and chromatin compaction, and gene expression (Alenghat and Ingber 2002; Dahl et al. 2008; Navarro et al. 2016).

There are two possible, ultimate (i.e., evolutionary) explanations for the consistent plastic phenotypic differences between cells cultured on soft and stiff substrates. First, it may be that the phenotypic outcome is optimal, i.e., there is no better way for a cell to perform its function on a given substrate. The tremendous morphological, physiological, and biochemical diversity of cell types in an organism derived from a common genotype suggests that this possibility cannot be summarily dismissed. Alternatively, it may be that the observed plasticity represents a compromise imposed by genetic and/or developmental constraints, such that improving the performance of a cell on one substrate necessarily reduces its performance on another. For example, if there is a temporal gradient in substrate rigidity over the course of development, optimizing cells for performance early may result in a phenotype that is constrained with respect to performance later, because the optimum late phenotype is developmentally inaccessible from the optimum early phenotype. That evolutionary optimization problem is analogous to that of life-history evolution, in which the age or stage that is most sensitive to changes in growth rate will constrain evolution at other ages/stages (Lande 1982; van Tienderen 1995; Williams 1957).

A promising way in which to uncover evolutionary constraints is by means of experimental evolution (Teotónio et al. 2017). For example, if optimal cellular performance on soft substrate is constrained by the need for cells to perform well on stiff substrate, and *vice versa*, then populations of cells selected for high fitness on a soft substrate are expected to evolve traits that are sub-optimal on stiff substrate, and *vice versa*. Conversely, if cells have evolved to have the optimal phenotype on both substrates (conditioned on the global constraint imposed by having to be an animal cell), then experimental evolution would produce no average change in phenotype (**Fig. 1,** purple arrow, “perfect plasticity“).

**Fig. 1.**
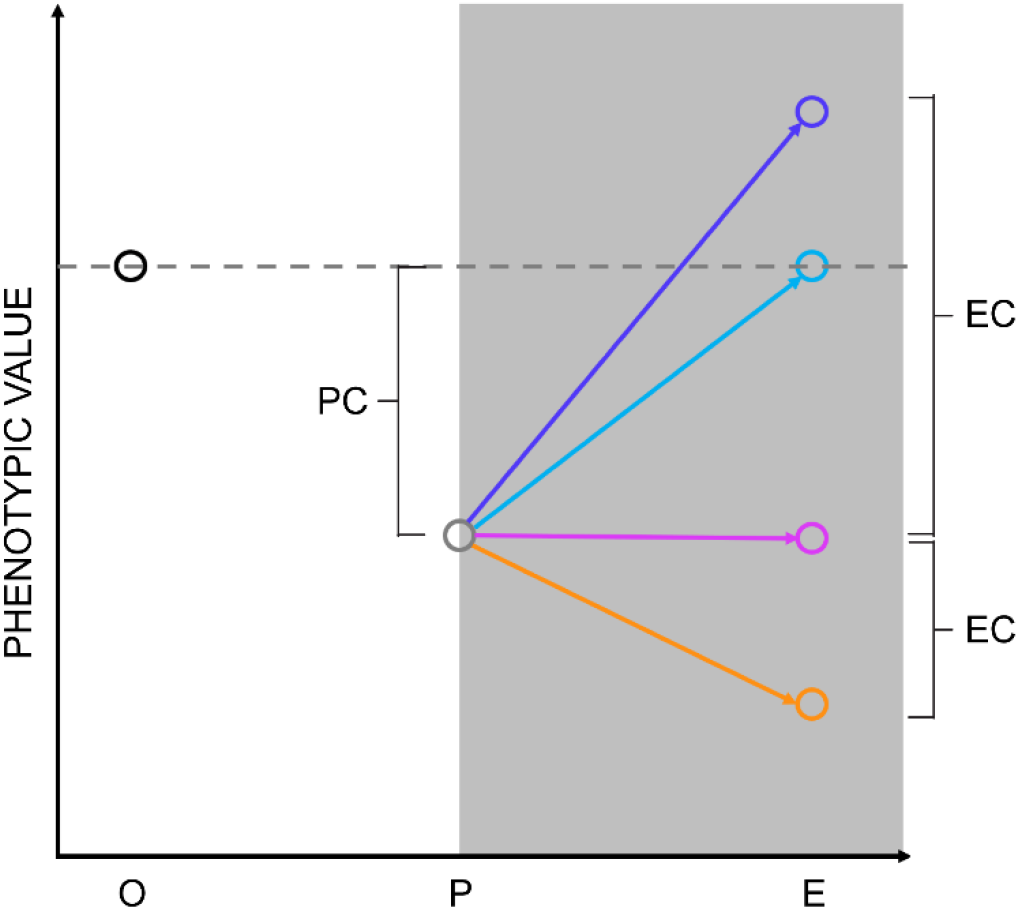
Correlation between plasticity and evolutionary outcomes. X-axis labels: O represents the phenotypic value of the ancestor on the stiff substrate; P represents the plastic phenotypic value of the ancestor on the soft substrate and E represents the phenotypic value after evolution on the soft substrate. PC designates “plastic change”, EC designates “evolutionary change” (see Results). The solid arrows represent potential trajectories of evolution; the dashed arrows represent the direction and magnitude of the constraint imposed by the ancestral fitness tradeoff.

Here we report a study in which we employ experimental evolution to characterize potential constraints on the phenotypic plasticity of mouse fibroblasts grown on soft and stiff substrates. We propagated replicate, genetically-variable populations of mouse fibroblasts, long adapted to a stiff substrate, for 90 cell-generations on soft (1 kPa) and stiff (308 kPa) substrates. To account for the possibility that phenotypic plasticity could manifest over longer time periods that might mimic genetic evolution, replicate clonal populations derived from single cells randomly sampled from the progenitor population were also propagated for 30 cell generations.

## Results

### Genetically variable populations of fibroblasts increased in fitness upon sustained culture on soft substrates

We chose as our source population, NIH 3T3 fibroblasts, which have been maintained in culture on a stiff substrate (plastic tissue culture dishes) for more than three decades and are a well-established model system for studying cellular sensitivity to substrate stiffness (Lo et al. 2000; Munevar et al. 2001b; Pelham and Wang 1997; Wang et al. 2014). The cells were initially derived from a highly inbred mouse and presumably were nearly completely homozygous, but have accumulated genetic variation for approximately 10^4^ generations (TODARO and GREEN 1963). Whole-exome sequencing of the source population revealed the fraction of segregating sites on the order of 1%. Of the 49,388,818 (mostly) exonic sites included in the final dataset, 746,629 (~1.5%) were segregating with a minor allele frequency (MAF) >0; 427,855 sites (~0.87%) had a MAF>1%.

Ten replicate lines were initiated by plating ~10^4^ cells on soft polyacrylamide gels (*E* = 1 kPa) and six replicate lines on stiff (*E* = 308 kPa) polyacrylamide gels (**Fig. 2A**). The specific values of stiffness were chosen because the plastic response of the cells on these two substrate stiffnesses has been previously reported (Lovett et al. 2013). Cell populations were allowed to grow for approximately three days and then 10^4^ cells were passaged onto fresh substrates of the same (corresponding) stiffness (**Fig. 2A;** cell doubling times were roughly 1-2 days). This procedure was carried out for three months, i.e., for approximately 90 cell generations. As a measure of fitness, we determined the cellular growth rate at one-month intervals over the course of the experiment (see methods).

**Fig. 2.**
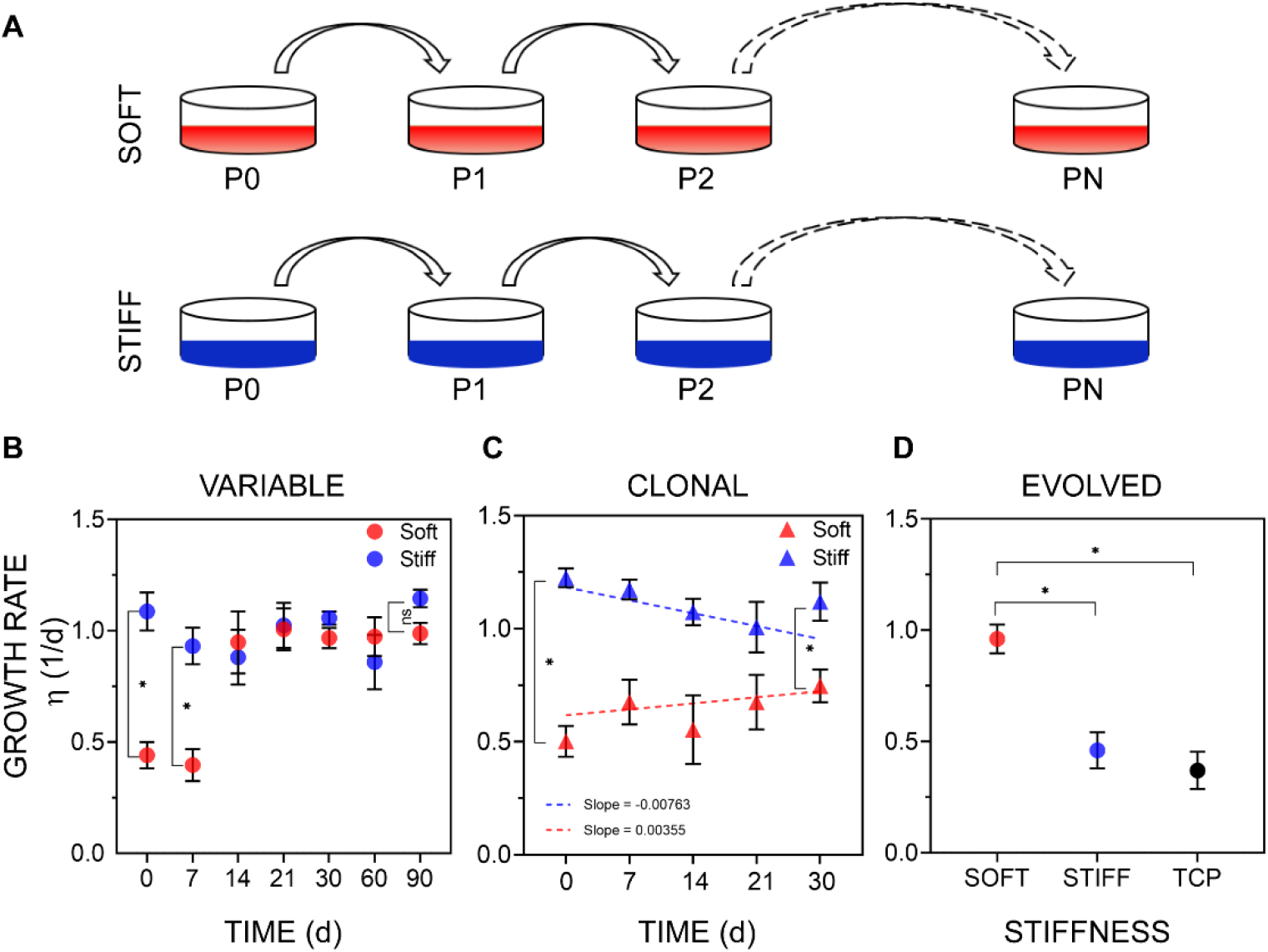
Evidence of evolution on soft substrates in murine fibroblasts. (A) Overview of the evolution experiment. Cells were cultured on the soft (red, E = 1 kPa) or stiff substrate (blue, E = 308 kPa) for up to 90 d. (B) The mean growth rate η (Units are 1/d; 1/η is the doubling time) of selected lines on the soft and stiff substrates, measured at different times during sustained 90-d culture. Error bars, SEM; data were collected from lines which were derived from the genetically variable ancestral population and were then cultured on the soft substrate (at least eight independent lines), or the stiff substrate (at least six independent lines). Statistically significant differences between treatments at each time-point were determined by Mann-Whitney U test, *P < 0.05; ns: P > 0.05. (C) The mean growth rate of lines derived from clonal cells determined at different times during sustained culture. Error bars, SEM; data were collected from eight or more different clonal lines. Difference between groups in the trajectory over the course of the experiment was assessed by General Linear Model (GLM; see Methods); also see supplemental **Fig S1**; regression slopes did not differ significantly from 0 (soft substrate, P>0.4, stiff substrate, P>0.06). (D) The mean growth rate of soft-selected lines on stiff substrates. Four lines were derived from the variable ancestral population and then cultured for 90 d on the soft substrate. These soft-selected lines were then cultured on soft or stiff substrates (TCP, tissue culture plastic) and their growth rate was measured on these substrates over 3 days of culture. Error bars, SEM. Statistical significance was determined by Kruskal-Wallis test followed Dunn’s multiple comparison test; *P < 0.05.

Consistent with previous studies (Mih et al. 2011), the initial growth rate of cells on the soft substrate was approximately half that on the stiff substrate (**Fig. 2B**). After 30 days of culture on the soft substrate, the growth rate had approximately doubled and was indistinguishable from the growth rate on the stiff substrate for the remainder of the experiment (**Fig. 2B**). However, assaying growth at 30-day intervals provides no information about the trajectory of the increase in growth rate over the initial 30-day period. To establish the early dynamics, we repeated these experiments for nine replicate lines on the soft substrate and ten replicates on the stiff substrate and measured the growth rate at seven-day intervals. The growth rate of the soft-selected lines on the soft substrate increased rapidly in approximately 2-3 weeks of culture, becoming statistically indistinguishable from the growth rate of both the ancestor and the stiff-selected lines on the stiff substrate. This rapid adaptation of growth rate to substrate stiffness has also been reported in other cell types (Syed et al. 2017).

To establish the generality of the phenomenon of rapid evolution of growth rate on a novel soft substrate, we conducted a similar experiment with a different cell type, murine C2C12 myoblasts (Yaffe and Saxel 1977), which also have been maintained on a stiff substrate (tissue culture plastic) for decades and thus are also expected to be similarly genetically variable. The myoblast growth rate on the soft substrate (n=6 lines) was initially lower than that of the same cells on the stiff substrate (n=6 lines). Over the course of two months, the growth rate of the soft-selected lines on the soft substrate had again evolved to be statistically indistinguishable from the growth rate of both the ancestor and the stiff-selected lines on the stiff substrate (supplemental **Fig. S2**). These observations indicate that the fitness of genetically variable cell lines can and does respond over the course of tens of generations to selection imposed by a novel substrate.

The rapid time scale and remarkable consistency of the increase in growth rate led us to suspect that the response could be a different manifestation of phenotypic plasticity. We reasoned that if the underlying cause of the increase in growth rate was phenotypic plasticity, then genetically homogenous (i.e., clonal) populations of cells should respond in the same way as the genetically variable populations. Accordingly, we repeated these experiments with clonal fibroblast lines initiated from individual cells (see methods), which are expected to be genetically homogeneous. Growth rate in clonal soft-selected lines did not change significantly over the course of 4 weeks (**Fig. 2C**). Thus, a genetically variable source population is required for an increase in growth rate on the soft substrate, consistent with the interpretation that the change in growth rate is the result of genetic evolution.

Unless selection is extraordinarily strong (e.g., as in the case of antibiotic exposure leading to drug resistance in bacterial culture), one would not expect that detectable evolutionary changes would occur on the timescale of a few cell divisions. The observed rapid evolution of growth rate on the soft (novel) substrate in the variable but not the clonal lines, combined with the two-fold difference in ancestral growth rate between the two substrates imply that the genotypes of the selected cells are present at low frequency in the ancestral population. Furthermore, if the selected genotypes are indeed rare in the ancestral population, this also implies that these genotypes must grow slowly on the stiff substrate because the ancestor had been selected on plastic dishes, which are stiff substrates. If either of those conditions did not hold, there would not be a big difference in ancestral growth rate on the two substrates. Consistent with the second prediction, we found that soft-selected lines grew slower on the stiff substrate than on the soft substrate (**Fig. 2D**).

With respect to the first prediction (i.e., that the selected genotypes are rare in the ancestral population), since somatic cells do not recombine during growth and the population size is not small (*N* = approximately 10^4^ cells), we can employ deterministic selection theory applied to competing clones to infer the approximate initial frequency and fitness advantage of the positively-selected genotypes (Haldane 1927). After *t* generations, the frequency of a focal clone, *p_t_*, can be calculated from the equation 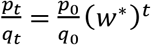, where *q* = 1–*p* represents the frequency of the competing clone(s); *p_0_* is the initial frequency of the focal clone; and *w*\* is the fitness advantage of the focal clone relative to wild type. The relationship between the initial relative frequency of a clone (*p_0_*), its relative fitness advantage (*w*\*), and its expected relative frequency after *t* generations (*p_t_*) is shown in supplemental **Table S1**. Comparison of the experimental values (supplemental **Fig. S3**, black squares) with the theoretical predictions revealed that the observations are consistent with a rapid increase in frequency (approximately 20 generations) of an initially rare clone with an approximately two-fold selective advantage over the wild type (*w*\*~ 2).

### Populations of cells evolved by natural selection and not random genetic drift

It is certain that genetically variable populations of cells will evolve over the course of 90 generations by random genetic drift even in the absence of selection. If the cause of the consistent change in growth rate is phenotypic plasticity, the only evolutionary force acting on the population would be random genetic drift (mutation notwithstanding). To investigate the possibility that the observed change in growth rate (and phenotype, see below) can be explained simply by random genetic drift, we performed whole exome sequencing of evolved and ancestral populations (see methods for details). Exomes of two sets of three pooled replicate soft-selected lines, one set of three pooled replicate stiff-selected lines, and a sample of the ancestral population were sequenced. We used the observed allele frequency spectrum of the ancestral population to parameterize simulations of *in silico* experimental evolution to establish the extent of sequence evolution that is consistent with selective neutrality. The allele frequency spectra were compared to each other and to the distributions simulated under the assumption of neutral evolution.

The distributions of allele frequencies in the four samples are shown in **Fig. 3A**. The thick black line in **Fig. 3A** shows the distribution of 500 neutral simulations. An example of the evolution of allele frequencies on a finer scale is shown in **Fig. 3B**. The distributions of the two sets of pooled soft-selected lines are very similar, and distinct from those of the pooled stiff-selected lines and of the ancestral population. When compared to the ancestor (purple curve in **Fig. 3A**), the frequency distribution of the soft-selected lines shows both a small peak of high-frequency variants (black arrow), as expected if some variants were increasing toward fixation, and a larger peak of low-frequency variants (blue arrow), as expected if a large fraction of variants was being removed from the population by selection. The same trend is evident in the stiff-selected sample, except that the peaks are at less extreme frequencies, which is consistent with directional selection being weaker (and hence the timescale of evolution slower) on the stiff substrate than on the soft substrate.

**Fig. 3.**
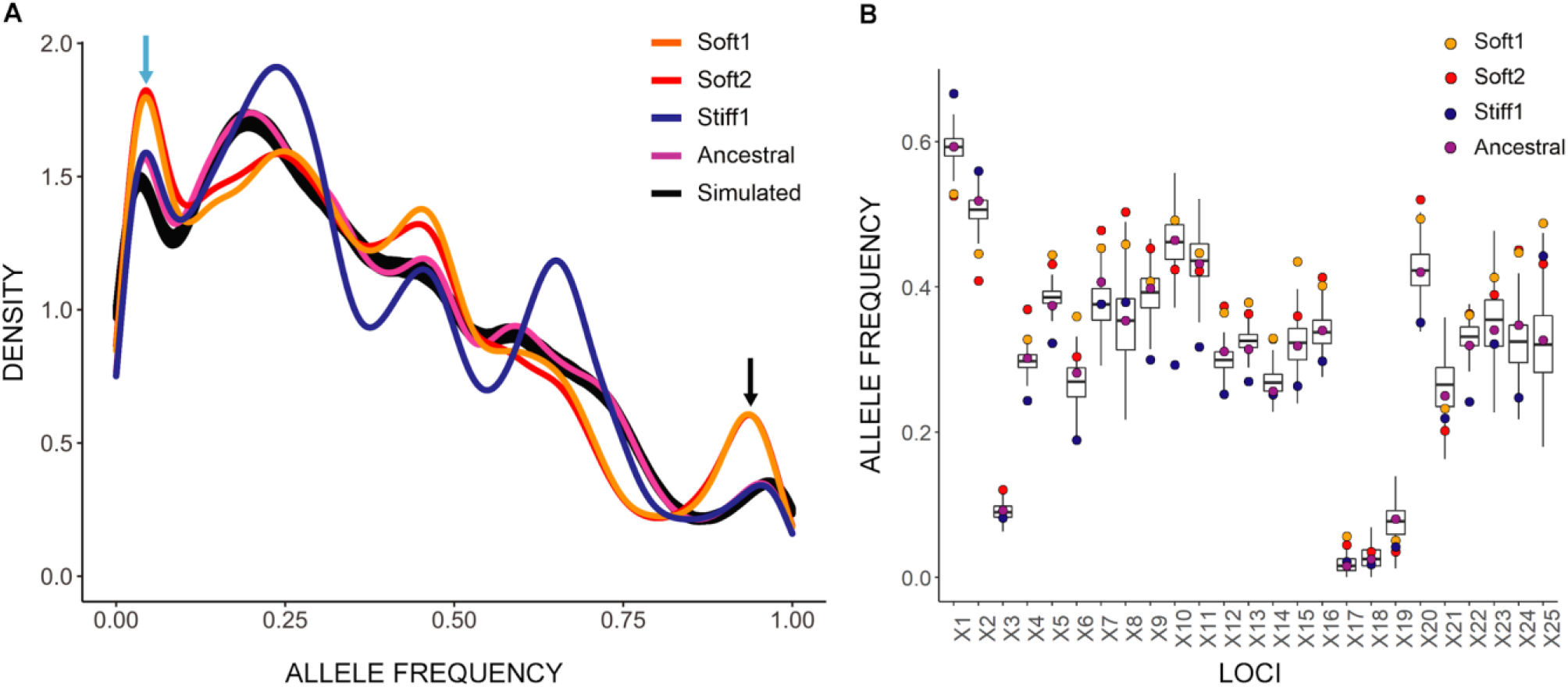
Density plot of alternative allele frequencies in the ancestral population and selected lines. (A) Two sets of three randomly chosen soft-selected lines were pooled to form two groups (Soft1 and Soft2). Three stiff-selected lines were also pooled (Stiff1). Exome sequencing of the pooled lines and ancestral population was performed at ~1000X average coverage, and allele frequencies calculated. The x-axis shows the unfolded allele frequency spectrum of variant alleles relative to the reference allele. The greater density of rare variants is presumably because the inbred mouse from which the cell line was derived was homozygous for the reference allele at most loci. The two pooled sets of soft-selected lines exhibit an excess of alleles that are close to fixation or extinction (black and blue arrows). Black line, allele frequencies calculated from 500 simulated trajectories in the absence of natural selection (see text). Change in allele frequency by locus, shown for the first 25 polymorphic loci on Chromosome 1. The boxplots represent the variation among 500 neutral simulations (see Methods for details of the simulations).

To quantify the extent of observed allele frequency change expected to occur in evolving populations as a result of random sampling (the cumulative contributions of random genetic drift, sample pooling, and sequencing), we simulated the 90 d sustained culture experiment described in **Fig. 2A** forward-in-time in the absence of selection and mutation. We constructed replicate lines as in the experiments *in silico* by randomly sampling 10^4^ completely linked diploid mouse genomes from a starting pool of 6 ⋅ 10^4^ genomes (each genome represents one cell) with the allele frequencies sampled with replacement from the observed allele frequency distribution in the ancestral population (purple curve in **Fig. 3A**). This simulated population was allowed to grow and evolve for 90 generations, bottlenecking to 10^4^ genomes at 3-generation intervals to represent cell passaging at three-day intervals. Two sets of three pooled lines were sequenced *in-silico* with the coverage mirroring the experimentally observed sequencing coverage at each locus.

As predicted by theory, the allele frequency changes over the course of 90 generations of neutral evolution in a population of N=10,000 are small. The average absolute per-site change in allele frequency (|Δ*p*|) across 500 simulations was 2.7% (simulation range 2.66% to 2.74%). The observed change in the experimental soft- and stiff-evolved lines was 4.4% and 3.7%, respectively (*P* < 0.002 based on 500 simulations). The average absolute relative change in allele frequency (|Δ*p*|/*p*) across 500 simulations was 14.1% (simulation range 13.8% to 14.4%). The observed change in the soft- and stiff-selected lines was 21.7% and 18.2%, respectively (*P* < 0.002). Further, 13.7% of sites in the soft-selected lines and 12.1% sites in the stiff-selected lines exceeded the maximum change for the sites across 500 simulations of neutral evolution. The observed changes in allele frequency over the course of 90 generations of evolution are too large to be explained by neutral evolution (plus sampling); the obvious conclusion is that the populations underwent adaptive evolution in response to natural selection imposed by the substrate.

To attempt to identify the causal alleles underlying adaptive evolution, we first filtered the exome-sequence data by the following criteria: (1) the minimum absolute allele frequency change |Δp|>0.0274 (the maximum value in 250 simulations), (2) minimum relative allele frequency change |Δ/pp|<0.141 (the maximum value in 250 simulations), (3) at least ten reads supporting the minor allele in at least one sample, and (4) the frequency of the minor allele in the ancestor less than 0.5. Of the 1394 variants that met those criteria (see supplemental excel sheet 1), we then looked for alleles that increased to a frequency of >25%, averaged over the two sets of soft-selected lines, remained at low frequency in the stiff-selected lines, and were covered >100X in at least one of the samples. Of the 19 variants that met those criteria, only one, a non-synonymous substitution in the carnitine transporter gene Slc22a2, reached high frequency (>95%) in both sets of soft-selected lines. However, both alleles were nearly absent in both the ancestor and the stiff-selected lines, suggestive of a false positive.

### Soft-selected cells interpret the soft substrate differently than ancestral cells

Evolutionary biologists typically make the *a priori* assumption that phenotypic plasticity points in the direction that natural selection favors (Price et al. 2003; Waddington 1942). For a hypothetical example, suppose a biannual plant reproduces twice a year, first in the wet season then in the dry season, and that broad leaves are favored when wet and narrow leaves favored when dry. Under those circumstances, natural selection would favor the evolution of a mechanism of phenotypic plasticity that produces broad leaves when wet and narrow leaves when dry. Now suppose a population of so-evolved plants finds itself in a consistently dry environment. We would predict that plasticity would (eventually) be lost, and that natural selection would produce a population with the same narrow leaves as the ancestor produced in the dry season (“perfect plasticity”, the purple line in **Fig. 1**). If the plasticity of leaf width in the ancestor was constrained by the selective requirement to produce the opposite leaf shape (broad v. narrow) in the opposite environment (wet v. dry), we would predict that the descendant population in the constant dry environment would eventually evolve leaves that were even narrower than those produced by phenotypic plasticity in the fluctuating ancestral environment (“evolutionary head-start”, the orange line in **Fig. 1**). However, there is strong empirical evidence that the phenotypic plasticity of certain kinds of traits – gene expression in particular – often points away from the direction of selection (blue lines in **Fig. 1**) (Ho and Zhang 2019) A caveat is in order, however, which is that most such examples are from systems in which the plastic response is initially measured in a novel environment, and the plastic response may represent a transient emergency response to environmental stress (Ghalambor et al. 2007).

We evaluated three traits in fibroblasts that have been previously shown to exhibit plasticity in cells cultured on substrates of different rigidities: cell spreading, Yes-associated protein (YAP) localization in the nucleus (Dupont et al. 2011), and cytoskeletal organization (Chan and Odde 2008). After three months of evolution on the soft substrate, both cell spreading area and nuclear/cytoplasmic ratio of YAP were statistically indistinguishable from the ancestral value on the stiff substrate (**Fig. 4A–D**). By contrast, these traits did not change over the course of three months in the stiff-selected lines. When cultured on soft substrates, ancestral cells do not assemble F-actin structures organized into linear contractile fibers and distinct microtubules, unlike ancestral cells cultured on the stiff substrate (**Fig. 4E**). Following evolution on the soft substrate, the cells gained the ability to assemble distinct F-actin fibers and microtubule networks comparable to ancestral cells on the stiff substrates. These results indicate that the mean phenotype tends to “revert to normal” on soft substrates (blue lines in **Fig. 1**), where normal refers to the state of the ancestor on stiff substrate. Just as for cell growth rate, phenotypic evolution required genetic variability in the starting population, because it was not observed in clonal cell populations (supplemental **Fig. S4**). The fact that genetically variable populations consistently evolved in the same way, and that genetically homogeneous clonal lines evolved far less (if at all) than the variable lines; **Fig. 2C**), strongly implies that the underlying cause of the change in mean phenotype in the genetically variable lines is the result of selection among genetically distinct clones, rather than phenotypic plasticity in a genetically uniform population manifested over a longer time scale.

**Fig. 4.**
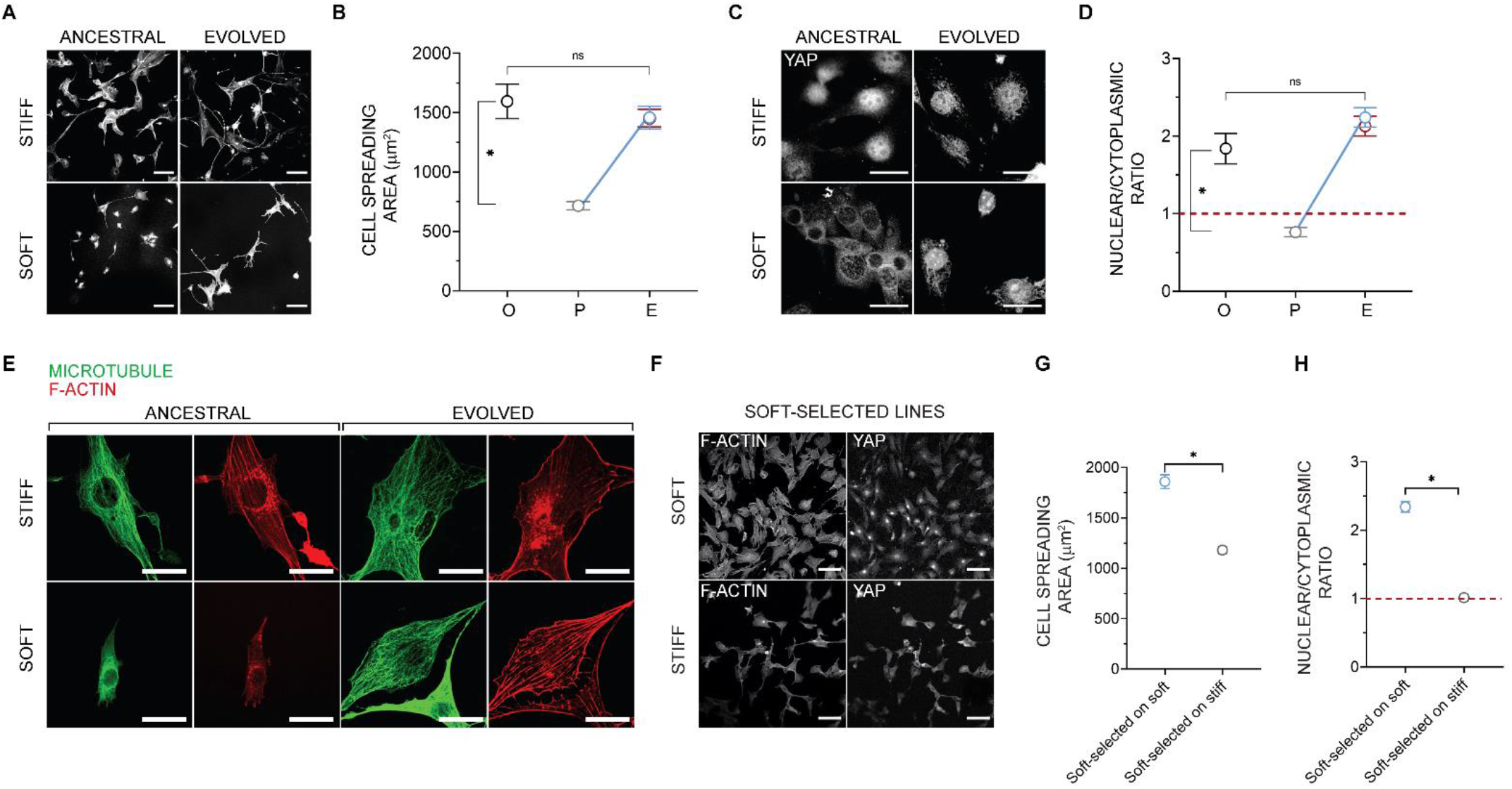
Phenotypic evolution. (A) Representative images of F-actin in ancestral populations cultured on the soft or stiff substrate for 3 d, and of evolved lines after 90 d of culture on the soft or stiff substrate; scale bar: 100 μm. (B) Mean spreading area of the ancestral cells on stiff substrate (O), ancestral cells on soft substrate (P) and selected lines after sustained culture (A) on soft (blue) or stiff substrates (red) (A); compare with the scheme in Figure 1. Error bars, SEM; each data point represents the mean value from over 50 cells from three lines (randomly chosen); Kruskal-Wallis test with Dunn’s multiple comparisons test, *P < 0.05; ns, P > 0.05. (C) Representative images of YAP in ancestral populations cultured on the soft or stiff substrate for 3 d, and in selected lines after 90 d of culture on the soft or stiff substrate; scale bar: 30 μm. (D) Mean nuclear to cytoplasmic YAP intensity ratio in ancestral cells on stiff substrate (O), ancestral cells on soft substrate (P) and selected lines after sustained culture (A) on soft (blue) or stiff substrate (red) (A); compare with the scheme in Figure 1. Error bars, SEM; each data point represents the mean value from over 50 cells from three lines; Kruskal-Wallis test with Dunn’s multiple comparisons test, *P < 0.05; ns, P > 0.05. (E) F-actin and microtubule structures in ancestral cells and selected lines after sustained culture on soft and stiff gels. Scale bar: 30 μm. (C and E). (F) Representative images of F-actin and YAP in soft-selected lines cultured on the soft or stiff substrate for 3 d. Scale bar: 100 μm (G) The mean spreading area of soft-selected lines on soft and stiff substrates. Error bars, SEM; each data point represents the mean value from over 225 cells from five or more lines (randomly chosen); Statistical significance was determined by Mann-Whitney U test; *P < 0.05. (H) The mean nuclear to cytoplasmic YAP intensity ratio of soft-selected lines on soft and stiff substrates. Error bars, SEM; each data point represents the mean value from over 200 cells from five or more lines (randomly chosen); Statistical significance was determined by Mann-Whitney U test; *P < 0.05.

Taken together, the results imply the following: in the ancestral population, which has been maintained for many generations on stiff substrate, there exist, at low frequency, cells with variant genotypes that (i) confer high growth rate on soft substrate, (ii) have a substantially lower growth rate on stiff substrate, but also (iii) manifest on the soft substrate the same suite of traits that the wild-type cells manifest on the stiff substrate. Apparently, there is a single optimal (multivariate) phenotype that confers high growth rate, and that phenotype is the same on stiff and soft substrates. This also suggests that the pattern of phenotypic plasticity exhibited by the selected cells must differ from the common (wild-type) pattern of plasticity present in the ancestral population. Otherwise, cells of the soft-selected lines grown on soft substrate would not have the characteristics of wild-type ancestral cells grown on stiff substrate. Confirming this intuition, we found that the pattern of plasticity present in the soft-selected lines was the opposite of the ancestral pattern. That is, soft-evolved cells spread less on the stiff substrate than on the soft substrate (**Fig. 4G**), and the nuclear/cytoplasmic ratio of YAP was greater on the soft than on the stiff substrate (**Fig. 4H**).

To characterize the (presumably genetic) variation in phenotypic plasticity, we isolated nine fibroblast clones from the ancestral population, expanded them into clonal lines, and then cultured these clonal cells on soft and stiff substrates. The clonal cells had low areas on soft substrates, and high areas on stiff substrates, as expected, and the correlation was large and positive (Pearson’s r=0.87, supplemental **Fig. S5)**.

### Different replicate lines evolved the same novel phenotypes through distinct patterns of gene expression

Cells of the soft-selected lines grow at the same rate on the soft substrate as wild-type ancestral cells grow on the stiff substrate, and they develop the same suite of phenotypic traits on the soft substrate that wild-type ancestral cells develop on the stiff substrate. An obvious possible point of control is at the level of gene expression. We therefore used RNA sequencing to investigate differences in gene expression between soft-selected lines and the ancestor in populations of cells cultured on soft and stiff substrates. We used principal component analysis (PCA) to cluster all samples based on the respective gene expression profiles (**Fig. 5A**). PC1, which explains 51% of the variance, clearly separates the ancestral and soft-selected populations, irrespective of substrate stiffness. PC2, which explains 28% of the variance, separates the soft-selected populations into two distinct clusters, but again there is no correspondence between the clusters and substrate stiffness. The lack of correspondence between clusters and substrate stiffness indicates that the soft-selected lines in the two clusters evolved distinct patterns of gene expression, independent of substrate. Collectively, these observations suggest that different soft-selected lines are composed of at least two distinct cell genotypes, which achieve high fitness and the same suite of phenotypic traits via at least two distinct transcriptional architectures.

**Fig. 5.**
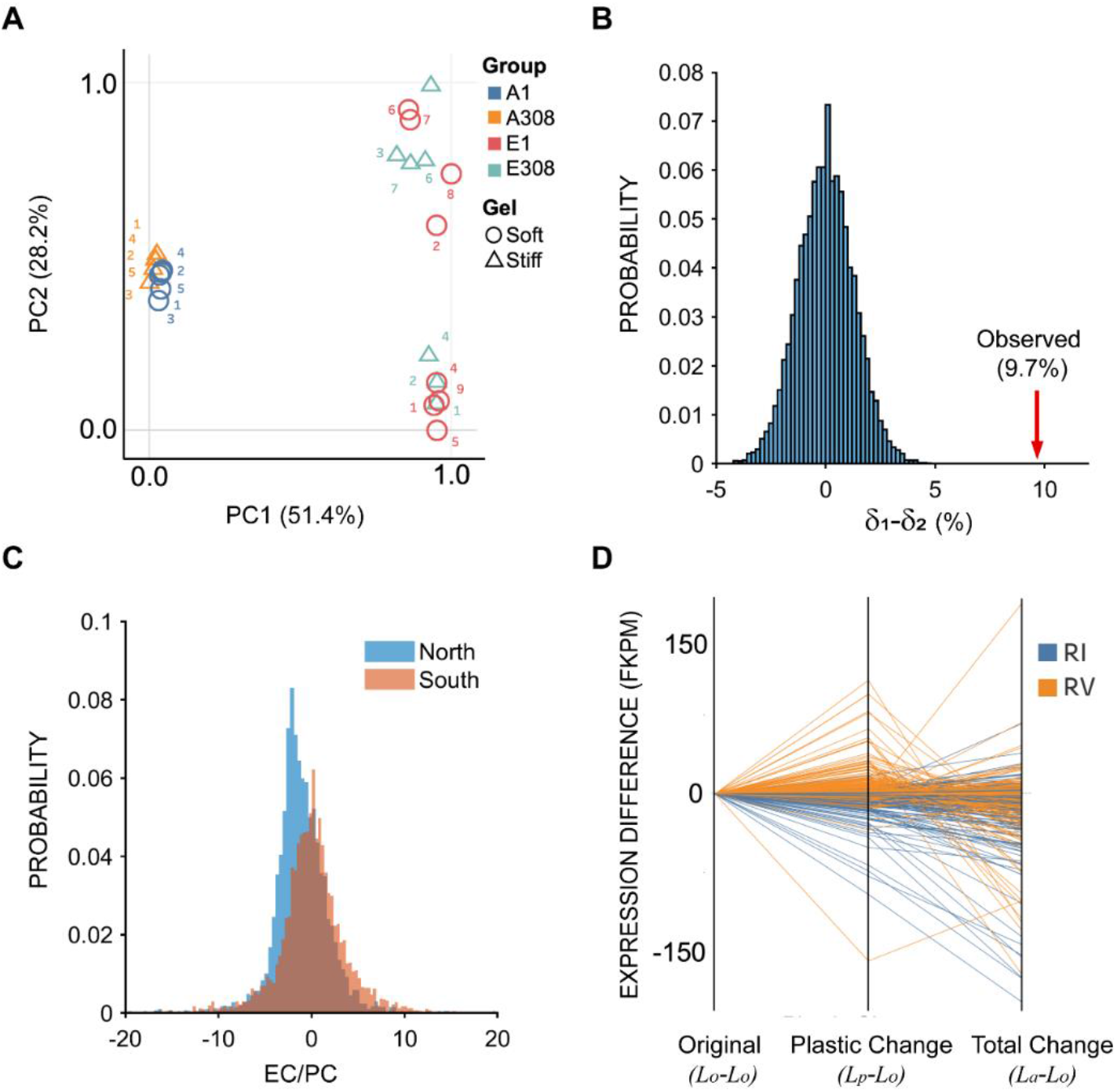
Analysis of gene expression. (A) Principal component analysis (PCA) of gene expression profiles. A1 and A308 represent ancestral cells cultured for 3 d on the soft (1 kPa) and stiff (308 kPa) substrates respectively. E1 and E308 represent cells evolved for 90 on the soft substrate (1 kPa) followed by 3 d culture on the soft (1 kPa) and stiff substrate (308 kPa) respectively. (B) Null δ_1_ –δ_2_ distribution generated by randomly sampling eight replicates from the *EC* and *PC* distributions (see text for definition of *EC* and *PC*). The actual value for δ_1_ –δ_2_ is indicated by a red arrow (see text). (C) Probability distributions of 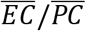 over all genes that exhibited statistically significant plasticity in the North (top cluster along PC2 in Fig. 5A) and South (bottom cluster along PC2 in Fig. 5A) clusters computed from parametric bootstrap simulations. (D) Parallel coordinate graphs depicting reversion or reinforcement of 449 genes identified as significantly reverted or reinforced from the parametric bootstrap simulation. Orange lines depict reversion (RV), blue lines depict reinforcement (RI); the indicated quantities at the bottom of the x-axis are plotted for each individual, identified gene.

The “revert to normal” pattern of phenotypic evolution observed on the soft substrate (**Fig. 4**) is consistent with similar patterns observed in both experimental (Fong et al. 2005) and natural populations (Ghalambor et al. 2015), in which gene expression has been shown to tend to revert to normal. Consequently, we asked whether gene expression also tends to revert to normal in the soft-selected lines. However, there is a known statistical bias for gene expression to apparently revert to normal (reversion) as opposed to becoming even more extreme (reinforcement) (Mallard et al. 2018). To account for this potential bias, we used a parametric bootstrap approach (Ho and Zhang 2019) to characterize all genes that exhibited statistically significant plasticity in the ancestor. This method involves re-sampling from a simulated normal distribution with the mean and variance equal to the observed values for each transcript (see methods). Using the sampled values, we calculated the plastic change (*PC*) and evolutionary change (*EC*; note that our EC is conceptually equivalent to the “genetic change”, GC, of Ho and Zhang 2018) as follows: *PC* = *L_p_* – *L_o_*, where *L_o_* and *L_p_* represent gene expression levels of the ancestor on the stiff substrate (the original environment) and upon plastic change (soft substrate); *EC* = *L_a_* – *L_p_*, where *L_a_* represents gene expression of the soft-selected lines on the soft substrate. The subscripts O, P, and A signify “original”, “plastic”, and “adapted”, respectively, following the terminology of Ho and Zhang (Ho and Zhang 2018). The ratio *EC/PC* captures both the magnitude and direction of change in gene expression upon evolution. If the ratio is negative, the gene expression is reverted (blue lines in Fig. 1), and if it is positive, the gene expression is reinforced (orange line in Fig. 1). The propensity of gene expression levels to revert or reinforce is captured by the parameter δ; δ = *C_RV_* – *C_RI_*, where *C_RV_* is the fraction of significantly reverted genes and *C_RI_* is the fraction of significantly reinforced genes. We found that δ for the eight soft-selected lines on the soft substrate was approximately 3.8%, indicative of a minor bias toward reversion (*P* < 0.05).

Next, to determine whether the difference along PC2 in the two clusters is associated with a difference in tendency to revert (RV) or reinforce (RI), we repeated the above analysis separately for the two clusters of evolved populations separated along PC2. For the North cluster, δ_1_ = 12.09%, indicating a trend toward reversion; for the South cluster, δ_2_ = 2.40% indicating a minor trend toward reversion. To address the possibility that the δ_1_–δ_2_ difference (9.7%) could be simply a result of sampling variance, we generated a null distribution of δ_1_ –δ_2_ by randomly sampling eight replicates from the distributions of *GC* and *PC*, and assigning them arbitrarily to one of two groups. The simulated null distribution is shown in **Fig. 5B**. The observed difference falls well outside the right tail of the distribution (red arrow in **Fig. 5B**), implying that the observed difference cannot be explained by sampling. These observations demonstrate that the separation of the two clusters along PC2 is caused at least in part by a difference between the two groups in the tendency of genes to revert. That is also apparent from the 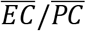 distributions over all genes that exhibit significant plasticity in the two groups (**Fig. 5C**). Specifically, the distribution is skewed toward negative values (i.e., reversion) for the North cluster, with no such trend in the South cluster. Fig. 5D provides a visual representation of reversion and reinforcement of individual genes which were identified from the parametric bootstrap simulation as significantly reverted or reinforced. Collectively (**Fig. 5B–D**), these observations indicate that different populations can evolve the same phenotypes by means of qualitatively different transcriptional pathways.

Because δ was significantly positive (indicative of a bias toward reversion) only for the North cluster, we performed gene ontology (GO) analysis of genes that reverted in the North cluster. These genes were enriched in biological processes related to mitosis and the cell cycle, and included microtubule-chromosome association (kinetochore organization), spindle organization, chromosome segregation, and regulation of cell cycle phases (supplemental **Fig. S6**; see methods). This is consistent with the phenotype returning to normal in terms of the growth rate on the soft substrate upon evolution (**Fig. 2**). It also suggests that the “revert to normal” of cell spreading and YAP nuclear localization likely regulate the expression of genes associated with growth rate.

### Gene expression changes in proliferative and mechano-transduction signaling networks

The preceding analyses shed light on the average properties of gene expression in the soft-selected lines, and on the general properties of the evolution of gene expression. However, they provide no information with respect to the specific genes and pathways that directly underlie the differences in substrate-specific growth rate and phenotypic plasticity between wild-type cells and the selected variants. Because cells sense and transduce mechanical stimuli into phenotypes and growth rates through the mechano-transduction pathway, and because evolved cells interpret the soft substrate differently from the ancestor, we set out to examine mechano-transduction pathways and growth pathways in lines evolved on the soft substrate. We specifically focused on the expression of genes in these pathways as measured by mRNA sequencing. These genes include the epidermal growth factor (EGF) pathway, the mitogen-activated protein kinase (MAPK) pathway, the Phosphoinositide-3 kinase (PI3K) pathway, and the mechano-transduction pathway. The genes involved in each pathway were obtained from the Kyoto encyclopedia of genes and genomes (KEGG) database (Kanehisa and Goto 2000) and further manually curated to remove overlap between pathways. The genes involved in the MAPK pathway were obtained from (Shi et al. 2016). The gene sets are provided in supplemental **Table S2**. To avoid confounding interpretations, we only included genes that are expected to be positively correlated with growth. Specifically, we did not include the negative regulators PTEN (PI3K) and Cofilin and GAP190 (mechano-transduction) in the analysis.

Using the t-test and an FDR of 10%, we identified genes that were differentially up- or down- regulated between the North and South clusters, each relative to the ancestral population. Given the baseline frequency *p* of up- or down- regulation estimated from all genes, we expect, by pure chance, that some genes in any pathway comprising *N* genes will be up- or down- regulated. We evaluated the statistical significance of observing up- or down- regulation of *k* or more genes in any pathway using the binomial distribution. The quantity 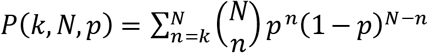 quantifies the probability that *k* or more genes are upregulated or downregulated in a pathway comprising *N* genes by pure chance. The probabilities can be found in supplemental **Tables S3 to S6**. Two distinct strategies appear to be employed by the south and the north clusters, respectively. The south cluster significantly upregulates the MAPK and the mechano-transduction pathways and to some extent the PI3K pathway. In contrast, the north cluster does not show any significant upregulation but shows significant downregulation in all considered pathways. We performed the same analysis for a manually curated set of genes known to be downstream of YAP (Wang et al. 2018). Similar to the analyses of growth factor and mechano-transduction pathways, we found that the YAP downstream targets were significantly upregulated in the south cluster and significantly downregulated in the north cluster. Consistent with the PCA above, this suggests mechanistically different evolutionary trajectories of the south and the north clusters. Specifically, the south cluster appears to evolve by increasing the strength of its coupling with the mechanical environment and the north cluster evolves by decoupling the growth machinery from external mechanical inputs.

## Discussion

Many animal cell types exhibit canonical phenotypic plasticity when cultured on substrates of differing stiffness. In this paper, we started with the premise that the mechanical properties of substrate are an inherent agent of natural selection for cells. We asked two questions: 1) First, do populations of cells respond to selection when continuously cultured on substrates of differing stiffness? 2) If cells do respond to selection, what type of phenotypic plasticity, gene expression patterns and sequence evolution are observed? If populations of cells do not respond to selection, that would imply either that there was no heritable variance for cellular fitness (at least on the relevant substrates), or that the cells were at evolutionary equilibrium on each substrate, consistent with the canonical plasticity resulting in the optimal cellular phenotype on each substrate.

We found that genetically variable populations of cells do respond to selection, but clonal populations do not. At the outset of the experiment, all populations of cells grew approximately twice as fast on stiff substrate as on soft substrate. As expected, they exhibited the canonical pattern of phenotypic plasticity with respect to cell spreading area, YAP localization, and cytoskeletal organization. Upon sustained culture on either soft or stiff substrate, genetically variable populations of cells cultured on stiff substrate maintained the ancestral growth rate, whereas growth rate in genetically variable populations of cells cultured on soft substrate increased to equal the growth rate on stiff substrate. In contrast, growth rate of genetically homogeneous populations of cells maintained under the same conditions did not change significantly. These results imply that evolution resulted from selection among existing genotypes (clones). Importantly, after 90 days of culture on the soft substrate, the soft-selected lines grew approximately half as fast when cultured on stiff substrate. Taken together, these results imply that the response to selection was driven by the increase in frequency of clones that were initially rare in the population that “did the wrong thing”, i.e., they grew fast on soft substrate and slowly on stiff substrate.

Strikingly, the soft-selected clones also exhibited the “wrong” pattern of phenotypic plasticity. That is, the pattern of cell spreading, YAP localization and F-actin assembly was reversed in the soft-selected lines (e.g., they spread more on soft substrate than on stiff substrate) from that of the ancestral lines. Clearly, the cells are not at a global optimum with respect to the two different substrate stiffnesses investigated here.

The observation that soft-selected cells “did the wrong thing”, i.e., growth rate and phenotypic plasticity were both reversed from the pattern in the ancestor, implies that the selected genotypes transduce the signal from the soft substrate in such a way as to produce the same suite of phenotypes that wild-type cells produce on the stiff substrate, and *vice versa*. That cells which evolved on soft substrates, spread less and also grew slower on the stiff substrate compared to the soft substrate, is consistent with the well-known direct relationship between cell spreading and growth rate (Chen et al. 1997). It is important to note that the observed reverse phenotypic plasticity of soft-selected cells (**Fig. 6B**) was not a foregone conclusion, and in fact was unforeseen. It is entirely conceivable that cells exist in which the (multivariate) phenotype is simply more extreme overall, in which case the plastic phenotype of soft-selected cells cultured on stiff substrate would be even more extreme than the ancestral (presumably optimum) mean (**Fig. 6C**), with a concomitant reduction in growth rate.

**Fig. 6.**
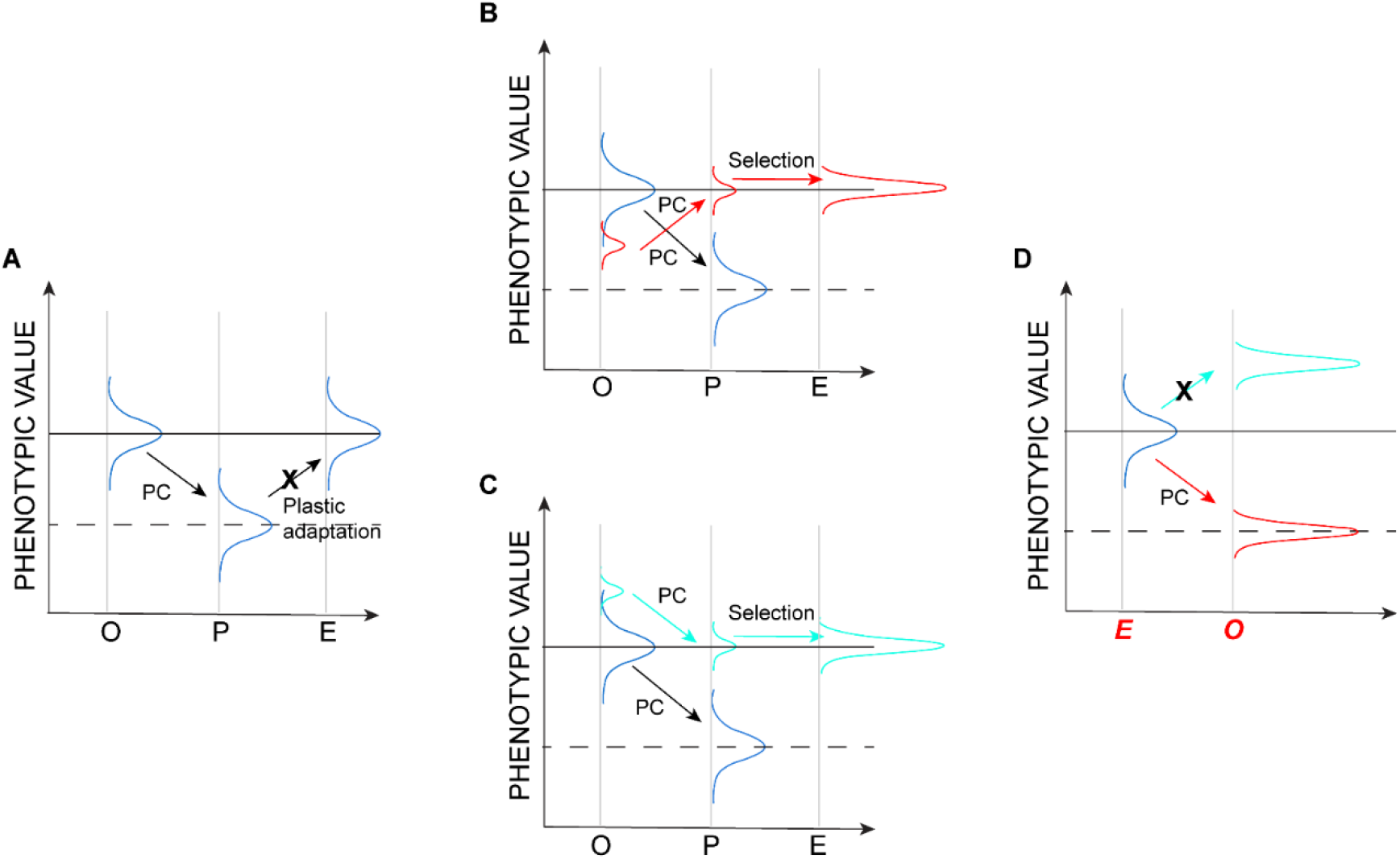
Plasticity and evolution revisited. Schematics show possible changes in hypothetical phenotypic distributions; labels on the X-axis in panels A-C are as in Figure 1. Arrows represent the direction of plastic change (PC). (**A**) The phenotypic response is the result of a change in phenotypic plasticity (“plastic adaptation“) over the long term. This scenario is ruled out (designated by an X) by the lack of observed change in clonal populations. (**B**) The evolutionary response results from selection of a rare clone (red) that “does the wrong thing”, i.e., its plastic response is opposite of the common types in the population. (**C)** The evolutionary response results from selection of a clone in the extreme upper tail of the phenotypic distribution (cyan) with a plastic response in the same direction as the common types in the population. (**D**) Predicted plasticity when evolved populations are exposed to the original environment. The scenario depicted in (B) predicts the observed outcome, shown in red; the scenario depicted in (C) predicts the outcome not observed, depicted in cyan.

Although the response to selection was remarkably consistent among replicates of the soft-selected lines, analysis of the transcriptome revealed underlying heterogeneity in the soft-selected lines. The first principal component, which explains half the variance in gene expression, clearly demarcates the soft-selected lines from the ancestor, irrespective of the substrate on which the samples were cultured. Thus, there is a common transcriptional response to selection imposed by substrate stiffness. PC2, which explains an additional 28% of the variance, separates the soft-selected lines into two groups, “North” and “South”, again irrespective of the substrate upon which the samples were cultured. Targeted analyses revealed differences between the North and South clusters in the expression of genes in the mechano-sensory pathway and genes involved in cell proliferation and growth. It appears that evolution on the soft substrate proceeded via one of the two alternatives; (1) by strongly coupling signaling with the environment (South) or (2) by decoupling signaling from the environmental input (North).

Our findings also inform a more general topic in evolutionary biology, concerning the relationship between phenotypic plasticity and adaptive evolution. As noted above, it is intuitive to think that phenotypic plasticity represents a stepping-stone on the way to further adaptation (“evolutionary head-start”, orange line in Figure 1), but it is well-understood that phenotypic plasticity does not always operate in that way, and that plastic phenotypes may often represent an emergency response to stressful circumstances (Ho and Zhang 2019). Although the NIH 3T3 fibroblasts have in fact been cultured on stiff substrates for thousands of cell generations, and in that context the soft substrate may legitimately be considered “novel”, animal cells have experienced variable substrate stiffness during the course of organismal development for hundreds of millions of years. From that perspective, it is not unreasonable to think that the common plastic response of cells to substrates of varying stiffness may in fact represent optimal phenotypic plasticity. Consistent with recent findings (Ho and Zhang 2019), our results argue against that interpretation, both at the level of emergent phenotypes and of gene expression, both of which tend to “revert to normal” in the soft-selected lines.

Ultimately, we would like to identify the specific genetic variants that are the targets of selection imposed by the mechanical environment. The results of the pooled exome sequencing revealed only one rather unconvincing variant that increased to high frequency in both (pooled) samples. Approximately 20 initially rare variants increased to ~25% in each pooled sample, although there were no obvious functional candidates among them. Because we sequenced exomes rather than whole genomes, it is conceivable (and we think likely) that the causal variants are regulatory rather than coding, and thus not represented in the exome sequence. A more fundamental limitation is that, because animal somatic cells do not recombine, all loci in the genome are completely linked, and population-level genome sequencing cannot unambiguously associate alleles with traits. An experiment in which individual cells are barcoded and sequenced at the single-cell level will, in principle, permit resolution down to the level of individual selected haplotypes, which could then be investigated based on understanding of the functions of the genes involved.

## Materials and Methods

### Experimental cell evolution on soft and stiff substrates

#### Synthesis and functionalization of hydrogels

Polyacrylamide gels of different stiffness were prepared by using a well-established protocol (Munevar et al. 2001a). Acrylamide and bis-acrylamide (Bio-rad) were mixed in 50:1 and 12.5:1 ratios to prepare gels with Young’s modulus (*E*) of 1 kPa and 308 kPa, respectively, as previously described (Lovett et al. 2013). We have previously shown that the cells sense gel stiffness in this assay and not other properties like ligand spacing (Lovett et al. 2013). The gel solutions contained 99.4% v/v gel mixture, 0.5% v/v ammonium persulphate (ThermoFisher Scientific), and 0.1% v/v tetramethylethylenediamine (ThermoFisher Scientific). For each gel, 100 μl of the gel solution was sandwiched between a hydrophobic glass surface and a hydrophilic glass coverslip (18 mm in diameter) for 20 min. The gels were functionalized using sulfosuccinimidyl 6-(4’-azido-2’-nitophenylamino) hexanoate (G Biosciences) and coated with fibronectin (10 μg/ml) before cell seeding.

#### Cell culture

The mouse fibroblast cell line NIH 3T3 and the mouse myoblast cell line C2C12 were purchased from the American Type Culture Collection. Both cell lines have been cultured for decades on (stiff) tissue culture plastic, and are expected to be at or near genetic equilibrium. The cells were cultured in 89% Dulbecco’s modified Eagle’s medium with 4.5 g/l glucose, 4 mM L-glutamine, and 1 mM sodium pyruvate (Corning Inc.), supplemented with 10% donor bovine serum (Gibco) and 1% penicillin-streptomycin mix (Corning Inc.). The cultures were maintained at 37 °C in a humidifier, under 7% CO_2_. The cells were detached from tissue culture dishes or gel surfaces using 0.25% trypsin (Corning Inc.) when the culture reached 70–80% confluence and were seeded onto new dishes/gels.

#### Cloning

NIH 3T3 fibroblasts were seeded on a dish (10 cm in diameter) at a density of 500 cells/ 10 cm diameter dish and were allowed to grow for 5–7 d. At that time, the cells formed 5–10 distinct colonies separate from one other. Each colony was isolated by placing a glass cloning ring (8 mm in diameter) on the surface of the tissue culture dish that enclosed a colony and trypsinizing the area of the dish enclosed by the cloning ring (McFarland 2000). The colonies were then grown independently in separate tissue culture dishes to obtain clones.

#### Growth rate measurements

Lines propagated on a given substrate were sampled at specific time points for growth rate measurements. Cells were trypsinized from the substrate of a given stiffness, a fixed number of cells were seeded on another substrate of the same stiffness, and allowed to grow for 3 d. On day 3, the cells were trypsinized from the gel surface. The trypsinized cells were suspended in 1 ml of the cell culture medium. Then, 10 μl of cell suspension was pipetted and mixed with 0.4% trypan blue (ThermoFisher Scientific) in 1:1 ratio. The cells were counted using Countess II Automated Cell Counter (ThermoFisher Scientific). The total live cell count/ml was recorded as the final number of cells and the growth rate was calculated as *N_f_* = *N*_0_ 2*^ηt^*, where *N_f_* is the final cell count; *N*_0_ is the initial cell count which was determined by the same procedure before seeding and was fixed at 10,000 cells; *t* is time (3 d); and *η* is the growth rate (per d).

#### Immunofluorescence staining and microscopy

Cells grown on the soft and stiff gels were fixed in 4% (wt/vol) paraformaldehyde (Alfa-aesar) in water, permeabilized with 0.2% Triton X-100 (Sigma) in PBS, and blocked with 1 mg/ml of bovine serum albumin (Sigma). Primary antibodies toward tubulin or YAP were added, and samples were incubated overnight at 4 °C. Subsequently, the samples were incubated with secondary antibodies for 2 h at room temperature. Microtubules were labeled using rabbit polyclonal anti-alpha tubulin antibody (ab18251; Abcam; dilution 1:600); and YAP was labeled using mouse monoclonal anti-YAP antibody (sc-271134; Santa Cruz Biotechnology; dilution 1:250). The secondary antibodies used were Alexa fluor 488-conjugated goat anti-rabbit antibody (A11008; Invitrogen; dilution 1:1000) and Alexa fluor 594-conjugated goat anti-mouse antibody (A11005; Invitrogen; dilution 1:1000). Nuclei were stained using Hoechst 33342 (Life Technologies) and F-actin was stained using Alexa fluor 647-conjugated phalloidin (A22287, Invitrogen). The gels were mounted on fluoro-dishes using ProLong Gold Antifade Mountant (ThermoFisher Scientific). Samples were imaged using a Nikon A1+ confocal microscope equipped with DU4 detector, and 20⋅ air or 60⋅ 1.4 NA oil-immersion objectives. Images were acquired using the NIS Elements 5.02 software (Nikon).

#### Cell area analysis

For cell area measurements, bright field images of cells attached to the gels were acquired using Zoe Fluorescent Cell Imager (BioRad); images of F-actin (following phalloidin staining in cells) were acquired using 20⋅ objective on a Nikon A1+ confocal microscope. Images were acquired at different locations on the gel surface and the area of each cell was determined using ImageJ software.

#### Measurement of the nuclear to cytoplasmic YAP ratio

The nuclear and cytoplasmic intensity of YAP was measured using ImageJ software. The nuclear boundary was traced in the fluorescent image, and the average YAP intensity in that area was measured using an ImageJ tool. The cytoplasmic intensity and background intensity were similarly determined by tracing areas. The nuclear to cytoplasmic YAP ratio was calculated as: [(Nuclear YAP intensity) – (Background intensity)] / [(Cytoplasmic intensity) – (Background intensity)]

#### Statistical analysis

The normality of distributions of the measured values was tested using the likelihood ratio test. The majority of distributions were not normal. Therefore, statistical comparisons between the control and treatment samples were done using the non-parametric Mann Whitney U test (Fig. 2B, 2C and Fig. S1). For multiple comparisons, Kruskal-Wallis test with Dunn’s multiple comparisons test was used (Fig. 2D, 4A, 4C). Graphical plots were generated using Prism version 8.4.0 (GraphPad), MATLAB version R2019b (Mathworks), and RStudio version 1.1.463 (RStudio Inc.). PCA data visualization (Fig. 5A, 5D) was done using R version 2.12.0 and Tableau Desktop version 2020.1. The details of experimental conditions and statistical tests are provided in figure legends.

The C2C12 growth rate data were analyzed by using the general linear model (GLM) implemented in the MIXED procedure of SAS version 9.4. The GLM can be written as

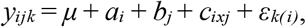

where *y_ijk_* is the measured growth rate of a sample; *μ* is the overall mean; *a_i_* is the continuous, linear effect of assay day *i*; *b_j_* is the fixed effect of substrate *j*; *c_ixj_* is the effect of the interaction between the assay day and substrate; and *ɛ_k(ij)_* is the residual effect. Variance components were determined using the restricted maximum likelihood method. The significance of fixed effects (day, substrate) was evaluated by F-test using Type III sums of squares. Degrees of freedom were determined by the method of Kenward and Roger (Kenward and Roger 1997). The linear regression of growth rate on assay day was then calculated separately for each substrate, using the linear model *y_ij_*=*μ* + *βt_j_*+ *ɛ_ij_*, where y*_ij_* is the growth rate of a sample; *μ* is the intercept and *β* is the slope of the linear regression of growth rate on assay day *t*; and *ɛ_ij_* is the residual effect.

### Whole-exome sequencing and analysis of gene expression in evolved lines

#### Whole-exome sequencing (WES) and variant analysis

To quantify evolution at the genomic level, WES (approximately 1000⋅ median coverage) was used to analyze a sample of the ancestral population, a pooled sample containing three fibroblast lines independently evolved for 90 days on the stiff substrate (308 kPa), and two pooled samples each containing three lines independently evolved for 90 days on the soft substrate (1 kPa). Whole exome was captured and sequenced using the Roche SeqCap® EZ MedExome Kit and Illumina NovaSeq^®^ 6000 Sequencer, respectively. To minimize sequencing errors, a sequencing protocol involving unique molecular identifiers was used (MacConaill et al. 2018) (xGen^®^ Dual Index UMI Adapters from Integrated DNA Technologies). Sequencing reads from each sample were mapped to a mouse reference genome version mm9 using BWA-MEM (Li and Durbin 2010). Library preparation, sequencing, de-multiplexing, read alignment, and quality check were performed according the standard laboratory protocols at the University of South California Genomics Core Facility. FreeBayes version 1.2.0 (Garrison and Marth 2012) joint variant calling method was used to deduce single-nucleotide variants, and small insertions and deletions in the aligned sequencing reads, yielding 750,873 variants. To maximize the capture of low-allele frequencies in the population, the following FreeBayes parameters were used: --pooled-continuous, --pooled-discrete, --min-alternate-fraction 0.001, --min-alternate-count 2, and --allele-balance-priors-off.

#### RNA extraction, library preparation, and sequencing

Five replicates of the ancestral population and 10 lines of fibroblasts evolved on soft 1 kPa gels for 90 d were each cultured on the soft or stiff gels for 3 d. After 3 d, each replicate population was trypsinized, centrifuged at 250 ⋅ *g* for 5 min, and counted to obtain at least 10^6^ cells per sample. Approximately 10^6^ cells were resuspended in RNAase-free PBS. The suspension was again centrifuged at 1400 ⋅ *g* for 3 min, the obtained pellet was flash-frozen in liquid nitrogen and immediately stored at –80 °C. The samples were shipped on dry ice to Novogene Corporation Inc.(Sacramento, California, USA) for RNA extraction and sequencing. There, the cell pellets were resuspended in RLT buffer supplemented with β-mercaptoethanol, lysed using lysis beads (Rodriguez-Palacios et al. 2015) and total RNA was extracted from the cell lysates using Qiagen RNAeasy mini kit according to the manufacturer’s protocol. The quality control of total extracted RNA was performed as follows: (1) preliminary quantitation using NanoDrop (ThermoFisher Scientific); (2) evaluation of contamination and RNA degradation by agarose gel electrophoresis; and (3) RNA integrity analysis using Agilent 2100 analyzer (Applied Biosystems). Samples containing over 0.8 μg of total RNA and with RNA integrity number > 7.0 were sequenced. Polyadenylated RNA was isolated using oligo(dT) beads. It was then randomly fragmented in a fragmentation buffer using enzyme method, and cDNA was synthesized using random hexamers and reverse transcriptase (included in the NEB Next® Ultra™ RNA Library Prep Kit for Illumina). After the first-strand synthesis, the second strand was synthesized using nick-translation in a customized second-strand synthesis buffer (Illumina) containing dNTPs, RNase H, and *Escherichia coli* polymerase I. The cDNA library was purified, and underwent terminal repair, A-tailing, ligation, size selection, and PCR enrichment, using NEB Next® Ultra™ RNA Library Prep Kit for Illumina (New England BioLabs). Library concentration was determined using Qubit 2.0 fluorometer (Life Technologies); the library was then diluted to 1 ng/μl, and insert sizes were checked using Agilent 2100 analyzer followed by quantitative PCR (Q-PCR) for quantifying to a higher accuracy(library activity>2nM). Paired-end 150-bp (PE150) sequencing (20 million raw reads) was performed using an Illumina HiSeq sequencer.

#### RNA sequencing and raw data analysis

Reads with adaptor contamination were trimmed and short reads (< 30 bp) were removed using Trimmomatic (Bolger et al. 2014). Cleaned reads were mapped to the mouse genome (version mm9) via HISAT2 (version 2.1.0) (Kim et al. 2015). The counts of aligned reads at the gene level were calculated using featureCounts (Subread version 1.6.2) (Liao et al. 2014) corresponding to the *Mus musculus* Ensembl 67 gene annotations. Weakly expressed genes were removed by retaining only the genes with at least 0.5 counts per million (CPM) in at least one sample. After the filtering step, 15,250 genes out of 35,275 genes remained, and were used in subsequent analysis.

For sample clustering and PCA analysis, counts data were transformed using the regularized log (rlog) method. PCA analysis and data visualization were performed in R using the pcaExplorer package (version 2.12.0) (Marini and Binder 2019) based on the top-1000 most variable genes across all samples.

Expression differences between ancestral cells cultured for 3 d on soft substrate (A1) or stiff substrate (A308), and cells pre-evolved on the soft substrate for 90 d and post cultured for 3 d on soft (E1) or stiff substrates (E308) were analyzed by using DEBrowser (version 1.14, DE method: DEseq2, test type Wald) (Kucukural et al. 2019). Adjusted *P*-value < 0.05 served as a cut-off to determine differentially expressed genes. Differential gene expression was visualized using iDEP.90 (Ge et al. 2018).

Functional categories over-represented in different subsets of genes were determined using WebGestalt 2019 (Liao et al. 2019) with the following parameters: enrichment method, over-representation analysis; minimum number of genes in a category, 20; FDR method, Bonferroni. Non-redundant GO biological processes with FDR < 0.05 are reported.

#### Parametric bootstrapping

Parametric bootstrapping was done as described elsewhere (Ho and Zhang 2019). For each transcript, random numbers were drawn from an assumed normal distribution with the observed mean and standard deviation calculated from the RNA sequencing data. Means and standard deviations were calculated for each transcript in the ancestral lines on the stiff and soft substrates, and in cells evolved on the soft substrate. Using these sampled values, *PC* (plastic change) and *EC* (evolutionary changes) were calculated; *PC* = *L_p_* – *L_o_*, where *L_o_* and *L_p_* represent gene expression levels in the original environment (stiff substrate) and upon plastic change (soft substrate), respectively; *EC* = *L_a_* – *L_p_*, where *L_a_* represents gene expression on the soft substrate after evolution. These calculations were repeated 10000 times for every transcript. If more than 9500 events of reversion, i.e., a change in sign from *EC* to *PC,* occurred, the transcript was classified as reverted. If the signs were the same, it was classified as reinforced. *C_RV_* was calculated as the fraction of genes that reverted, while *C_RI_* was calculated as the fraction of genes that reinforced.

### Forward-in-time simulations of experimental evolution

Evolution in a finite population is inevitable because of the random sampling of genomes, i.e., the genetic drift. Sampling variance in the sequencing coverage among loci and among samples will also lead to observed allele-frequency change. To establish the magnitude of allele-frequency change (Δ*p*) consistent with the cumulative effect of genetic drift and sampling of genomes during sequencing of the experimental populations, forward-in-time simulations of *in silico* populations were implemented with the same expected initial allele frequency spectra and demography as those of the actual experimental populations. The following genomic variants (see section on WES above) were excluded from analysis: 1. Those variants whose observed frequencies were consistently over 99% or under 1% across all samples, and 2. Those variants whose “Qual” score was less than 30. This resulted in 42,004 polymorphic genetic variants that were included in the simulation study.

The mouse cells studied here are diploid and genomes do not recombine. Each line was initiated by randomly sampling 10,000 cells (= 20K haploid genomes) from a pool of 60,000 cells with an allele frequency spectrum equivalent to that of the ancestral experimental population (see section on WES above). At each variable site in each simulated (haploid) genome, the allele was assigned with an expectation equal to the observed frequency *p_0_*. The sampling protocol in effect assumes that genomes are initially in linkage equilibrium, which is not true. However, because variable sites are sparse relative to the read length and because individual cells were not sequenced in the current study, the initial haplotype structure was not known. Neutral allele-frequency dynamics at individual loci do not depend on the assumption of linkage equilibrium.

Subsequent to initiation, each population of 10,000 cells went through three cycles of cell division, with the assumption that every cell divides (i.e., there is no cell death), to a confluent population size of 80,000 cells. This mimics the seeding densities and expected division rates of cells in the evolution experiment of Fig. 2A. Haploid genomes were paired in cells, and pairs of genomes doubled with each cell division event. From this population of 80,000 cells, 10,000 cells were randomly drawn without replacement for the next round of population growth, mimicking the passage of cells in the experiment at regular intervals (Fig. 2A). This sampling process was repeated for 26 cycles, nearly equivalent to *N* = 78 cycles of cell growth in the actual experiment. At the terminal generation, lines were randomly pooled into three “pool-seq” sets of three replicates to mimic the experimental exome sequencing approach. The allele-frequency, *p*′, was calculated from the entire sample of 60,000 alleles after sampling the corresponding number of reads for each locus.

The observed difference in allele frequency, *p*′ – *p_0_*, is an unbiased estimate of the change in allele frequency, Δ*p*, but it underestimates the sampling variance in Δ*p* because the observed allele frequency in the ancestor is an estimate rather than a known parameter. To re-estimate *p*′ after accounting for the additional sampling variance associated with the estimation of *p_0_*, we generated a sample of *n* binomially sampled alleles using *p*′ (as the probability of success), where *n* is the observed coverage at the locus in the ancestor. We call the mean of the *n* sampled alleles *p*\*, and the variance in *p*\* accounts for all sources of sampling variance: sequencing variance of the ancestor; random genetic drift over the course of the experiment; variation associated with the pooling of replicates of evolved populations; and sequencing variance of the pooled evolved lines. The simulation process was repeated 500 times.

Because the loci are completely linked, the upper bound of the uncorrected *P*-value of an observed Δ*p* greater than the most extreme simulated value is 1/500, and the lower boundary is the inverse of the product of the number of variable sites (*N* = 42,004) and the number of simulations, i.e., 1/(500*N).

## Acknowledgments

We thank the anonymous reviewers for their helpful suggestions, in particular Reviewer #2 for helping clarify our discussion of the relationship between plasticity and clonal selection, and for suggesting the depiction in Figure 6.

## Author contributions

Conceptualization, T.P.L., S.C. and C.F.B.; Methodology, Investigation, P.P., K.P., A.S.S. and H.H.; Software, A.S.S. and T.P.L. ; Formal Analysis, P.P., H.H., A.S.S., T.P.L., C.F.B. and S.C.; Writing-Original Draft, T.P.L., A.S.S.,C.F.B. and P.P.; Writing-Review and Editing, P.P., A.S.S., H.H., T.P.L. and C.F.B; Supervision, S.C., T.P.L. and C.F.B.

## Data and materials availability

All data is available in the main text or the supplementary materials and will be deposited into Dryad upon acceptance for publication. Codes used for parametric bootstrapping and for simulating evolution forward-in-time are available at https://github.com/srikarchamala/cellular_evolution_paper. All sequencing data is currently deposited at NCBI BioProject Accession ID PRJNA626453 and will be available to public as soon as the manuscript is published.

