## Supplemental Information for "Reverse plasticity underlies rapid evolution by clonal selection within populations of fibroblasts propagated on a novel soft substrate"

**This file includes:**

Figs. S1 to S6
Table S1 to S6

**Supplementary Figures:**

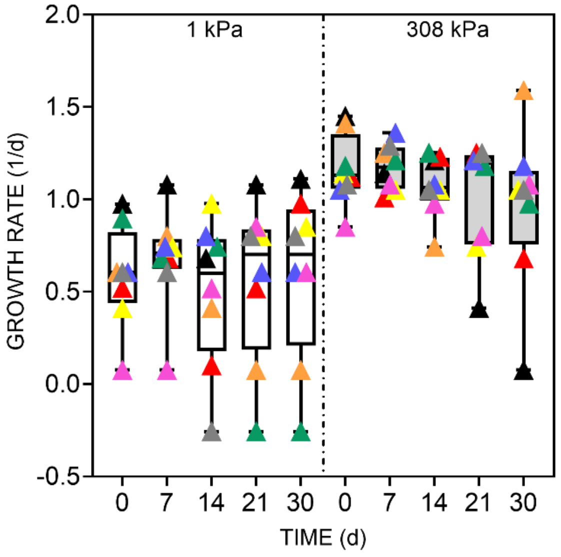


**Fig. S1. Growth trajectories of individual clones.** Growth rates of eight distinct clonal populations, measured at different times during their sustained culture on soft (1 kPa) or stiff (308 kPa) substrate for 30 d. Each color represents an individual clone; the color is the same for a given clone on gels of either stiffness.

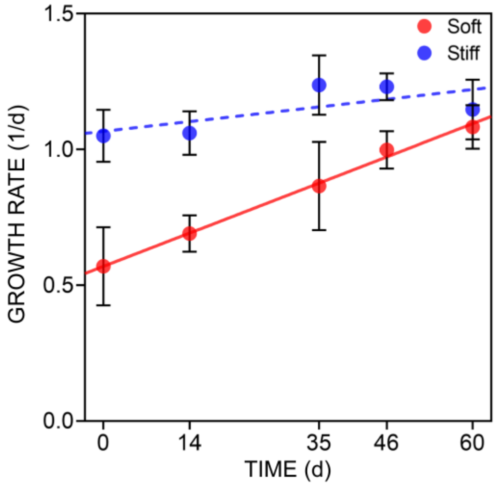


**Fig. S2. Measurements of growth rates in C2C12 myoblasts.** The mean growth rate of ancestral C2C12 myoblast cell populations on the soft and stiff substrates (1 kPa and 308 kPa respectively) was measured at different times during sustained culture over a 60-d period. Error bars, SEM; data were collected from five or more lines on the soft substrate, and six lines on the stiff substrate at each time point. Statistical significance of the overall trend was determined by GLM (see SI Methods). The lines represent the linear regression of growth rate in time (d). For the growth on the soft substrate (red continuous line; $P < 0.001$), slope = 0.008767 and y-intercept = 0.5694; for the growth on the stiff substrate (blue dashed line; $P > 0.18$), slope = 0.002566 and y-intercept = 1.067. GLM analysis revealed a significant linear increase in growth rate on the soft substrate, but no increasing trend in growth rate on the stiff substrate, as indicated by the significant *day* × *substrate* interaction in the linear model ($F_{1,50.2} = 4.69$, $P < 0.04$). The growth rate increased by approximately 0.88%/d on the soft substrate ($F_{1,25} = 16.24$, $P < 0.001$), whereas there was no significant change in the growth rate on the stiff substrate ($F_{1,27} = 1.87$, $P > 0.18$).

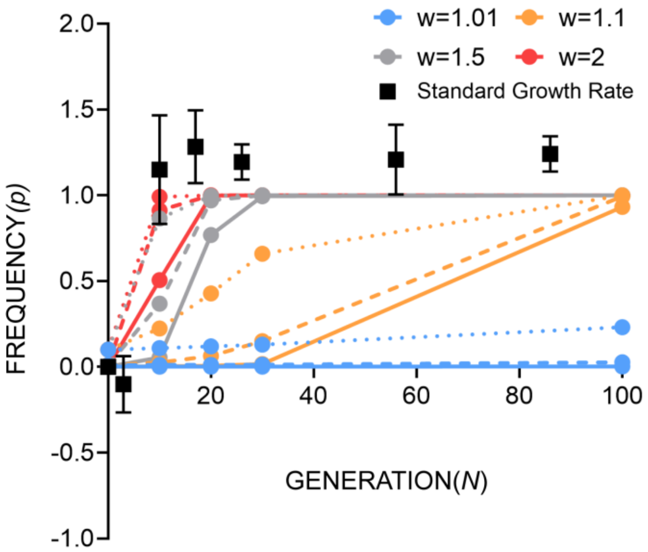


**Fig. S3. Fitness advantage of the focal clone relative to wild type.** The colored lines depict deterministic clone frequency dynamics from a given starting frequency $p_0$, with a given strength of selection *w*, representing the relative fitness of the focal clone relative to the population mean. Solid lines, $p_0 = 0.001$; dashed lines, $p_0 = 0.01$; and dotted lines, $p_0 = 0.1$. Colors denote the strength of selection, as defined in the figure key. Black squares represent the experimentally measured mean growth rate of selected fibroblast lines on the soft substrate (1 kPa) relative to the ancestor (note that the growth rate at generation 3 is lower than the ancestral value); errors bars are S.E.M. Ancestral value was scaled to 0; a value of 1 represents a 100% increase in growth rate.

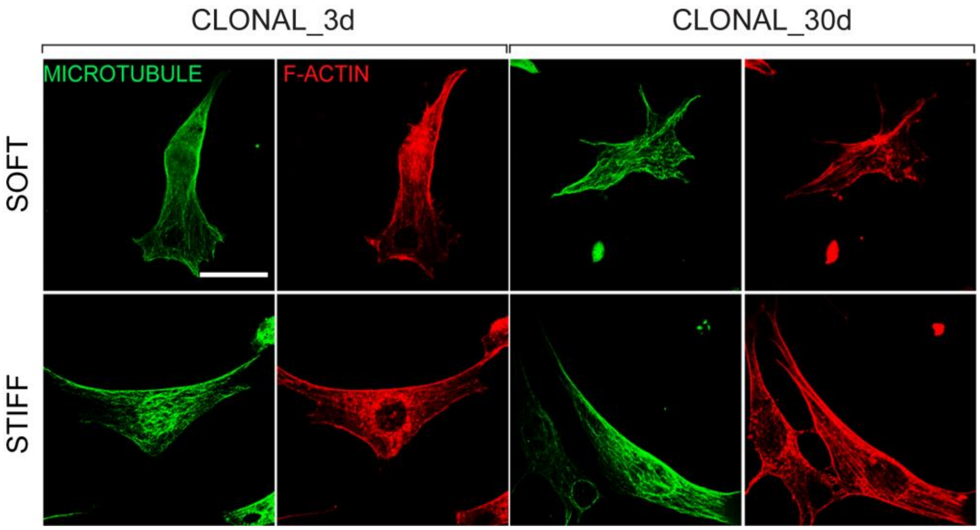


**Fig. S4. Cytoskeletal organization in clonal fibroblasts on the soft substrate.** Microtubule and F-actin structures in clonal fibroblasts before and after 30 d of sustained culture on the soft and stiff substrates. Scale bar: 30 μm. The cells were immunostained (see methods) and imaged with a 60x objective on a Nikon A1+ confocal microscope. The images are representative of over 100 cells of at least three clonal populations.

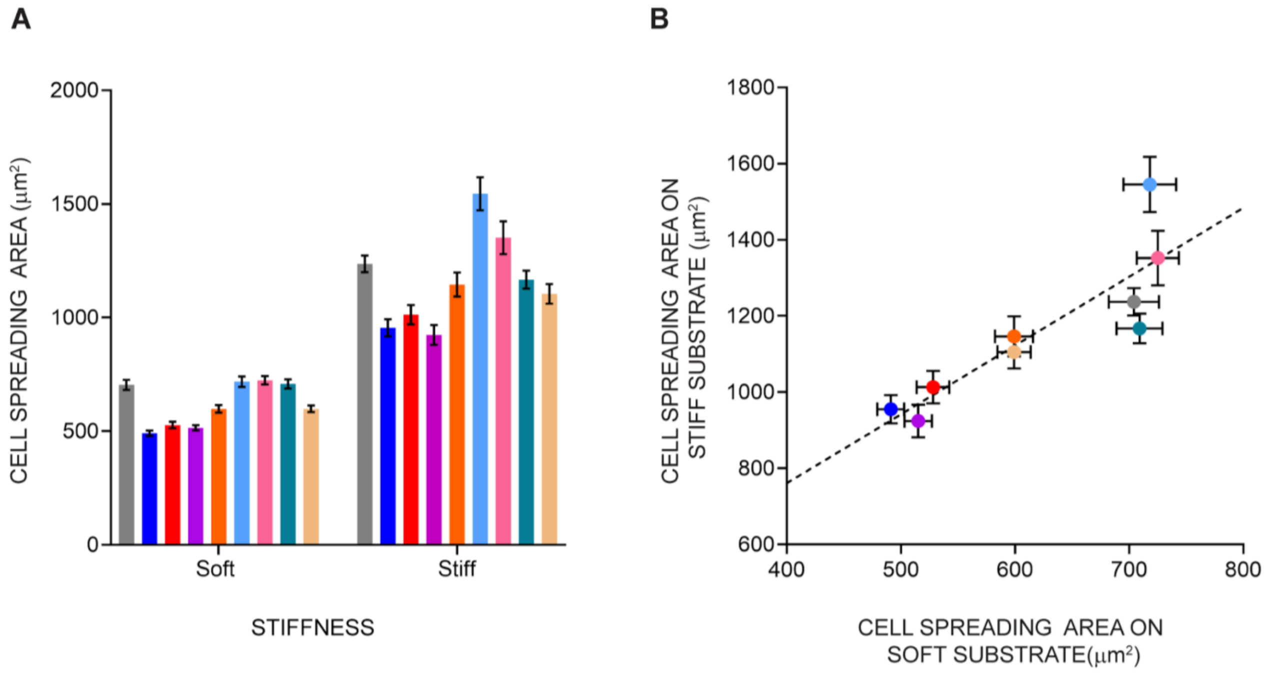


**Fig. S5. Area of cell spreading of clonal populations on soft and stiff substrates.** (A) Mean area of cell spreading of nine distinct clonal populations, each cultured on soft (1 kPa) or stiff (308 kPa) substrate for 3 d. Error bars, SEM; mean was calculated from at least 140 different cells from three different gels of a given stiffness for each clone. Each color represents an individual clone; this color is the same for a given clone on gels of either stiffness. (B) Mean spreading area of clonal populations from A) on stiff and soft substrates re-plotted against each other. Error bars are S.E.M. The color scheme is the same as in A. Pearson correlation coefficient, $r$ = 0.87.

A.

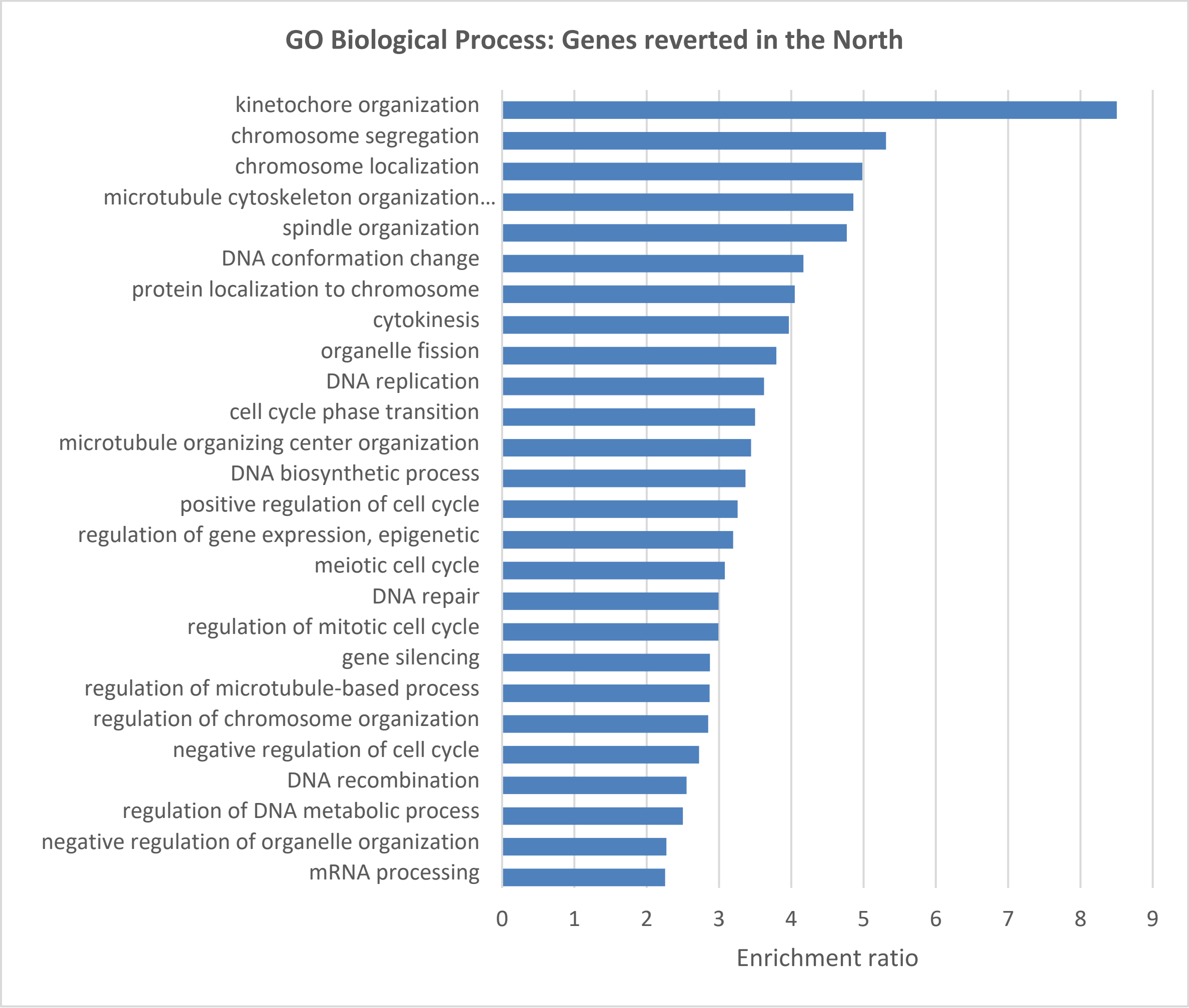
GO Biological Process: Genes reverted in the North
kinetochore organization
chromosome segregation
chromosome localization
microtubule cytoskeleton organization...
spindle organization
DNA conformation change
protein localization to chromosome
cytokinesis
organelle fission
DNA replication
cell cycle phase transition
microtubule organizing center organization
DNA biosynthetic process
positive regulation of cell cycle
regulation of gene expression, epigenetic
meiotic cell cycle
DNA repair
regulation of mitotic cell cycle
gene silencing
regulation of microtubule-based process
regulation of chromosome organization
negative regulation of cell cycle
DNA recombination
regulation of DNA metabolic process
negative regulation of organelle organization
mRNA processing
0 1 2 3 4 5 6 7 8 9
Enrichment ratio

B.

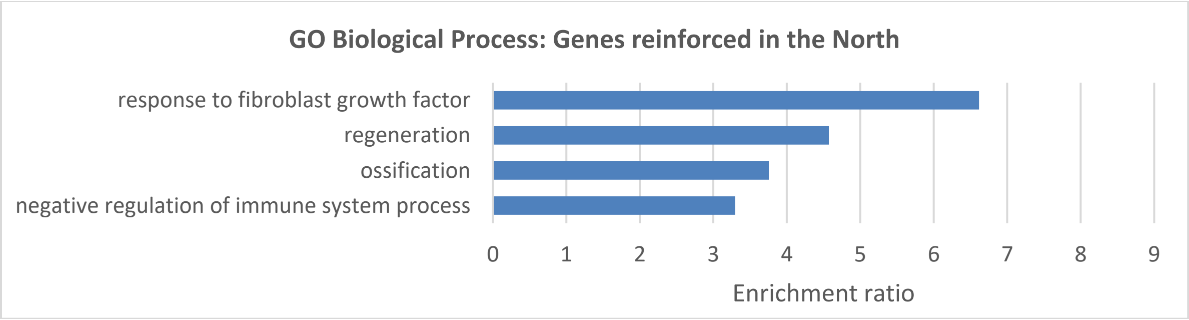
GO Biological Process: Genes reinforced in the North
response to fibroblast growth factor
regeneration
ossification
negative regulation of immune system process
0 1 2 3 4 5 6 7 8 9
Enrichment ratio

C.

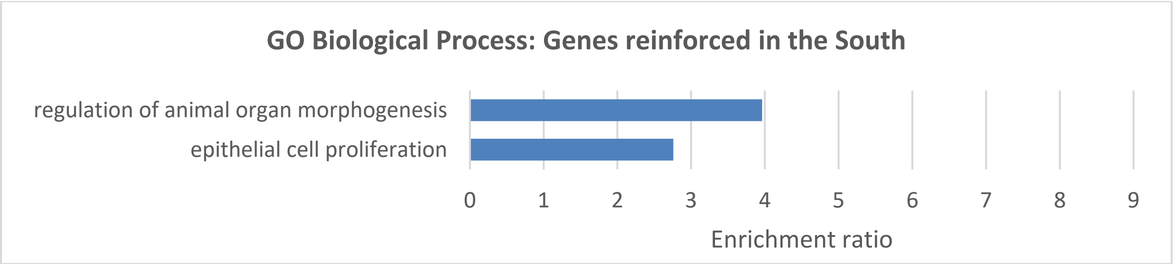


D.

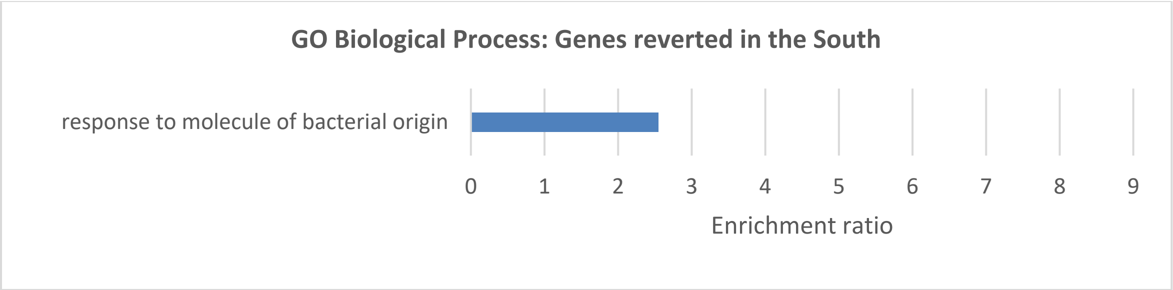


E.

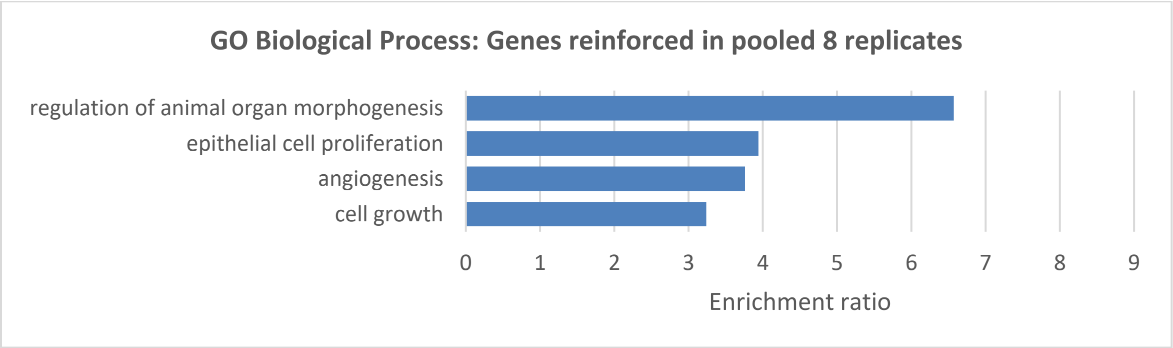


F.

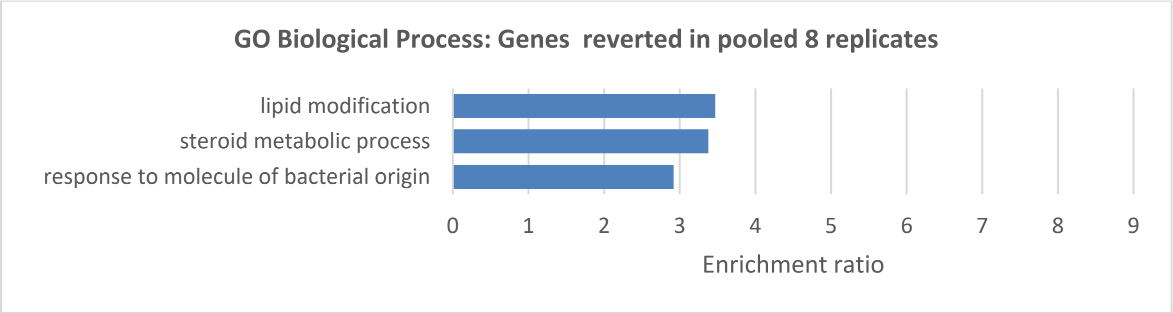


**Fig. S6. Gene ontology analysis of the reverted and reinforced genes.** The results of over-representation analysis to determine over-represented gene categories identified by parametric bootstrap simulations as reverted or reinforced. Comparisons were made between the gene set of interest and the reference transcriptome (Mus musculus genome protein coding). Results are shown for the North cluster (A and B; see Fig. 5A for nomenclature), South cluster (C and D), and all pooled samples (E and F; eight replicates evolved on the soft substrate).

**Supplementary Tables:**

| *p0* | *w*/w<sub>wt</sub>* | *p10* | *p30* | *p100* | *p300* |
|---|---|---|---|---|---|
| 0.0001 | 1.01 | 0.000 | 0.000 | 0.000 | 0.002 |
| | 1.05 | 0.000 | 0.000 | 0.013 | 0.996 |
| | 1.1 | 0.000 | 0.002 | 0.580 | 1 |
| | 1.25 | 0.001 | 0.075 | 1 | 1 |
| | 1.5 | 0.006 | 0.950 | 1 | 1 |
| | 2 | 0.093 | 1 | 1 | 1 |
| 0.010 | 1.01 | 0.011 | 0.013 | 0.027 | 0.167 |
| | 1.05 | 0.016 | 0.042 | 0.571 | 1 |
| | 1.1 | 0.026 | 0.150 | 0.993 | 1 |
| | 1.25 | 0.086 | 0.891 | 1 | 1 |
| | 1.5 | 0.368 | 0.999 | 1 | 1 |
| | 2 | 0.912 | 1 | 1 | 1 |

| *p0* | $w^*/w_{wt}$ | *w10* | *w30* | *w100* | *w300* |
|---|---|---|---|---|---|
| 0.0001 | 1.01 | 1.00 | 1.00 | 1.00 | 1.00 |
| | 1.05 | 1.00 | 1.00 | 1.00 | 1.05 |
| | 1.1 | 1.00 | 1.00 | 1.06 | 1.1 |
| | 1.25 | 1.00 | 1.02 | 1.25 | 1.25 |
| | 1.5 | 1.00 | 1.48 | 1.5 | 1.5 |
| | 2 | 1.09 | 2 | 2 | 2 |
| 0.010 | 1.01 | 1.00 | 1.00 | 1.00 | 1.00 |
| | 1.05 | 1.00 | 1.00 | 1.03 | 1.05 |
| | 1.1 | 1.00 | 1.01 | 1.10 | 1.1 |
| | 1.25 | 1.02 | 1.22 | 1.25 | 1.25 |
| | 1.5 | 1.18 | 1.50 | 1.5 | 1.5 |
| | 2 | 1.91 | 2 | 2 | 2 |

**Table S1**. **Relationship between the relative frequency and relative fitness of a competing clone.** The left panel represents deterministic clone frequency dynamics starting from the initial frequency $p_0$. $w^*$ is the growth rate of the focal clone; $w_{wt}$, the growth rate of the wild type scaled to 1 and p($t$), the frequency of the focal clone after $t$ generations. Gray background denotes the time at which $P > 0.5$. The right panel represents population mean growth rate after $t$ generations. A population composed of 100% wild-type cells has a growth rate of 1; a population composed of 100% focal clones has a growth rate of $w^*$. Gray background denotes the time at which the mean growth rate reached > 90% of its final value.

A. EGF pathway

| Gene name | Mouse gene ID |
|---|---|
| EGF | ENSMUSG0000002801 7 |
| TGF-alpha | ENSMUSG0000002999 9 |
| AREG | ENSMUSG0000002937 8 |
| BTC | ENSMUSG0000008236 1 |
| HBEGF | ENSMUSG0000002448 6 |
| EREG | ENSMUSG0000002937 7 |
| NRG1a | ENSMUSG0000006299 1 |
| NRG1b | ENSMUSG0000011854 1 |
| NRG2 | ENSMUSG0000006027 5 |
| NRG3 | ENSMUSG0000004101 4 |
| NRG4 | ENSMUSG0000003231 1 |
| EGFR | ENSMUSG0000002012 2 |
| ERBB2 | ENSMUSG0000006231 2 |
| ERBB3 | ENSMUSG0000001816 6 |
| ERBB4 | ENSMUSG0000006220 9 |

B. MAPK pathway

| Gene name | Mouse gene ID |
|---|---|
| ARAF | ENSMUSG00000001127 |
| BRAF | ENSMUSG00000002413 |
| CRAF | ENSMUSG00000000441 |
| GAB1 | ENSMUSG00000031714 |
| GRB2 | ENSMUSG00000059923 |
| HRAS | ENSMUSG00000025499 |

| | |
|---|---|
| KRAS | ENSMUSG00000030265 |
| MAP2K1 | ENSMUSG00000004936 |
| MAP2K2 | ENSMUSG00000035027 |
| MAPK1 | ENSMUSG00000063358 |
| MAPK3 | ENSMUSG00000063065 |
| NRAS | ENSMUSG00000027852 |
| RASA1 | ENSMUSG00000000441 |
| SHC1 | ENSMUSG00000021549 |
| SOS1 | ENSMUSG00000024241 |
| SOS2 | ENSMUSG00000034801 |

C. PI3K pathway

| Gene name | Mouse gene ID |
|---|---|
| Akt1 | ENSMUSG00000001729 |
| PDK1 | ENSMUSG00000024122 |
| PIK3R1 | ENSMUSG00000041417 |
| PIK3R2 | ENSMUSG00000031834 |
| PIK3R3 | ENSMUSG00000028698 |
| PIK3R4 | ENSMUSG00000032571 |
| PIK3R5 | ENSMUSG00000020901 |
| PIK3R6 | ENSMUSG00000046207 |
| PI3KCA | ENSMUSG00000027665 |
| PI3KCB | ENSMUSG00000032462 |
| PI3KCG | ENSMUSG00000020573 |
| PI3KCD | ENSMUSG00000039936 |
| MTOR | ENSMUSG00000028991 |
| S6K1 | ENSMUSG00000020516 |
| S6K2 | ENSMUSG00000024830 |

D. Mechanotransduction pathways

| Gene name | Mouse gene id |
|---|---|
| CCRK | ENSMUSG00000017776 |
| BCAR1 | ENSMUSG00000031955 |
| FAK | ENSMUSG00000022607 |
| YAP | ENSMUSG00000053110 |
| ROCK | ENSMUSG00000024290 |
| MDIA | ENSMUSG00000024456 |
| RHOA | ENSMUSG00000007815 |
| TEAD | ENSMUSG00000055320 |

| | |
|---|---|
| TAZ | ENSMUSG00000027803 |
| VINCULIN | ENSMUSG00000021823 |
| PAXILLIN | ENSMUSG00000029528 |
| ZYXIN | ENSMUSG00000029860 |
| GEF190 | ENSMUSG00000021662 |
| PIP4K2A | ENSMUSG00000026737 |
| PIP4K2B | ENSMUSG00000018547 |
| PIP4K2C | ENSMUSG00000025417 |
| DOC180 | ENSMUSG00000058325 |
| MYOSIN2 | ENSMUSG00000013936 |
| SRCK | ENSMUSG00000027646 |

E. Genes downstream of YAP/TAZ

| Gene name | Mouse gene id |
|---|---|
| CYR61 | ENSMUSG00000028195 |
| CTGF | ENSMUSG00000019997 |
| AMOTL2 | ENSMUSG00000032531 |
| ANKRD1 | ENSMUSG00000024803 |
| IGFBP3 | ENSMUSG00000020427 |
| F3 | ENSMUSG00000028128 |
| FJX1 | ENSMUSG00000075012 |
| NUAK2 | ENSMUSG00000009772 |
| LATS2 | ENSMUSG00000021959 |
| CRIM1 | ENSMUSG00000024074 |
| GADD45A | ENSMUSG00000036390 |
| TFGB2 | ENSMUSG00000039239 |
| PTPN14 | ENSMUSG00000026604 |
| NT5E | ENSMUSG00000032420 |
| FOXF | ENSMUSG00000038402 |
| AXL | ENSMUSG00000002602 |
| DOC5 | ENSMUSG00000044447 |
| ASAP1 | ENSMUSG00000022377 |
| RBMS3 | ENSMUSG00000039607 |
| MYOF | ENSMUSG00000048612 |
| ARHGEF17 | ENSMUSG00000032875 |
| CCDC80 | ENSMUSG00000022665 |

**Table S2**. **Gene IDs and names.** Gene sets involved in A.epidermal growth factor (EGF) pathway, B. mitogen-activated protein kinase (MAPK) pathway, C. Phosphoinositide-3 kinase (PI3K) pathway, D. mechanotransduction pathways and E. YAP/TAZ pathway.

North cluster upregulation

| Pathway | # of genes | upregulated | P+ |
|---|---|---|---|
| EGFR | 14 | 0 | 0.99 |
| PI3K | 15 | 0 | 0.99 |
| MAPK | 16 | 2 | 0.33 |
| Mechanotransduction | 19 | 0 | 0.99 |
| YAP/TAZ | 22 | 1 | 0.81 |

**Table S3. Upregulation of gene expression for growth and mechanotransduction related pathways in the north cluster.** The upregulated genes were determined using a FDR cutoff of 10%. The probability P+ represents the likelihood of observing upregulation of k or more genes by chance; k is the number of observed upregulated genes.

South cluster upregulation

| Pathway | # of genes | upregulated | P+ |
|---|---|---|---|
| EGFR | 14 | 3 | 0.48 |
| **PI3K** | **15** | **5** | **0.12** |
| **MAPK** | **16** | **7** | **0.016** |
| **Mechanotransduction** | **19** | **11** | **0.001** |
| **YAP/TAZ** | **22** | **11** | **7x10-4** |

**Table S4. Statistics of upregulation of gene expression for growth and mechanotransduction related pathways in the south cluster.** The upregulated genes were determined using a FDR cutoff of 10%. The probability P+ represents the likelihood of observing upregulation of k or more genes by chance; k is the number of observed upregulated genes.

North cluster downregulation

| Pathway | # of genes | downregulated | P- |
|---|---|---|---|
| **EGFR** | **14** | **5** | **0.04** |
| **PI3K** | **15** | **10** | **5.9x10-6** |
| **MAPK** | **16** | **6** | **0.02** |
| **Mechanotransduction** | **19** | **14** | **10-8** |
| **YAP/TAZ** | **22** | **10** | **5x10-4** |

**Table S5. Statistics of downregulation of gene expression for growth and mechanotransduction related pathways in the north cluster.** The downregulated genes were determined using a FDR cutoff of 10%. The probability P- represents the likelihood of observing downregulation of k or more genes by chance; k is the number of observed downregulated genes.

South cluster downregulation

| Pathway | # of genes | downregulated | P- |
|---|---|---|---|
| EGFR | 14 | 1 | 0.67 |
| PI3K | 15 | 0 | 0.99 |
| MAPK | 16 | 0 | 0.99 |
| Mechanotransduction | 19 | 3 | 0.17 |
| YAP/TAZ | 22 | 2 | 0.50 |

**Table S6. Statistics of downregulation of gene expression for growth and mechanotransduction related pathways in the south cluster.** The downregulated genes were determined using a FDR cutoff of 10%. The probability P- represents the likelihood of observing downregulation of k or more genes by chance; k is the number of observed downregulated genes.
